# Direct Nanopore Sequencing of Individual Full Length tRNA Strands

**DOI:** 10.1101/2021.04.26.441285

**Authors:** Niki Thomas, Vinay Poodari, Miten Jain, Hugh Olsen, Mark Akeson, Robin Abu-Shumays

## Abstract

We describe a method for direct tRNA sequencing using the Oxford Nanopore MinION. The principal technical advance is custom adapters that facilitate end-to-end sequencing of individual tRNA molecules. A second advance is a Nanopore sequencing pipeline optimized for tRNA. We tested this method using purified *E. coli* tRNA^fMet^, tRNA^Lys^, and tRNA^Phe^ samples. 76-to-92% of individual aligned tRNA sequence reads were full length. As proof of concept, we showed that Nanopore sequencing detected all 42 expected tRNA isoacceptors in total *E. coli* tRNA. Alignment-based comparison between the three purified tRNAs and their synthetic controls revealed systematic miscalls at or adjacent to the positions of known nucleotide modifications. Systematic miscalls were also observed proximal to known modifications in total *E. coli* tRNA alignments.

## INTRODUCTION

tRNA plays a central role in protein synthesis and has non-translational regulatory functions^1^. They adopt a cloverleaf secondary structure that typically includes four loops: the T loop, the variable loop, the anticodon loop, and the D loop. The tRNA cloverleaf structure further folds into an L shape important for binding and function in the ribosome^2^. Mature tRNA molecules typically contain a terminal, single-stranded 3′NCCA end. Over 90 unique ribonucleotide modifications are documented among all tRNAs^3^.

tRNA sequencing is typically performed using RNA-seq^4,5^. This method employs reverse transcription (RT) and sequencing by synthesis of cDNA products. Certain modified bases cause RT to stop or stall which in some methods is mitigated using demethylase treatments or thermostable group II intron reverse transcriptases (TGIRTs)^6^ to obtain full length reads. However, RNA-Seq cannot directly detect base modifications.

Nanopore RNA sequencing is a distinctly different technique that reads nucleotides directly without RT or amplification steps^7^. This permits detection of modified nucleotides as part of the sequencing process^8–12^. Anticipating the use of nanopore technology for tRNA sequencing, Smith et al. (2015)^13^ developed a strategy for unfolding and threading tRNA strands through alpha-hemolysin pores. To accomplish this, double-stranded adapters were annealed to the tRNA NCCA 3′ overhangs and then ligated to the 3′ and 5′ termini. Using non-catalytic phi29 DNA polymerase as a brake, *E. coli* tRNA^fMet^ and tRNA^Lys^ molecules were classified based on ionic current duration and amplitude for three segments along each strand. However, RNA nucleotide sequencing was not possible.

Here, we build on that prior work and implement Nanopore sequencing of individual, full length tRNA strands using the Oxford Nanopore Technologies (ONT) MinION.

## MATERIALS AND METHODS

### Enzymes

T4 RNA Ligase 2 (10,000 units/mL), T4 DNA Ligase (2,000,000 units/mL), T4 Polynucleotide kinase (10,000 units/mL), and corresponding buffer solutions, were purchased from New England Biolabs. Nuclease P1 (Sigma Aldrich), Antarctic phosphatase (5,000 units/mL) (New England Biolabs), Phosphodiesterase 1 (Thermo Fisher Scientific) were used to prepare samples for Liquid chromatography/mass spectrometry.

### Biological tRNAs

Purified biological tRNAs included *E. coli* tRNA ^fMet^ (Subriden RNA), tRNA ^Lys^ (Sigma-Aldrich) and tRNA^Phe^ (Sigma-Aldrich). *E. coli* total tRNA from strain MRE600 was obtained from Roche Pharmaceuticals.

### Synthetic Canonical tRNA

Synthetic canonical tRNAs were ordered from IDT or Dharmacon. 5′ phosphorylation was performed either during synthesis (for tRNA^fMet^) or following synthesis using T4 Polynucleotide kinase (NEB)(for tRNA^Lys^ and tRNA^Phe^) per manufacturer′s protocol. The sequences of these tRNAs are:

*E. coli* synthetic tRNA^fMet^ (initiator tRNA) 5′PCGCGGGGUGGAGCAGCCUGGUAGCUCGUCGGGCUCAUAACCCGAAGGUCGUCGGUUCAAAUCC GGCCCCCGCAACCA3′
*E. coli* synthetic tRNA^Lys^ 5′PGGGTCGTTAGCTCAGTTGGTAGAGCAGTTGACTTTTAATCAATTGGTCGCAGGTTCGAATCCTGCA CGACCCACCA3′
*E. coli* tRNA^Phe^ 5′PGCCCGGATAGCTCAGTCGGTAGAGCAGGGGATTGAAAATCCCCGTGTCCTTGGTTCGATTCCGAGT CCGGGCACCA3′

#### Splint adapter oligonucleotides

There were four distinct double-stranded splint adapters, each designed for one of the four different 3′ overhangs of *E. coli* tRNAs. Each was composed of two synthetic oligomers purchased from IDT. The oligonucleotide that ligates to the 3′ end of the tRNA was common to all splint adapters (Fig 1A, *i* blue strand). It was composed of six ribonucleotides followed by 24 DNA nucleotides. The 30 nt sequence for this oligonucleotide was: 5′P-rGrGrCrUrUrCTTCTTGCTCTTAGGTAGTAGGTTC-3′.

**Figure 1.**
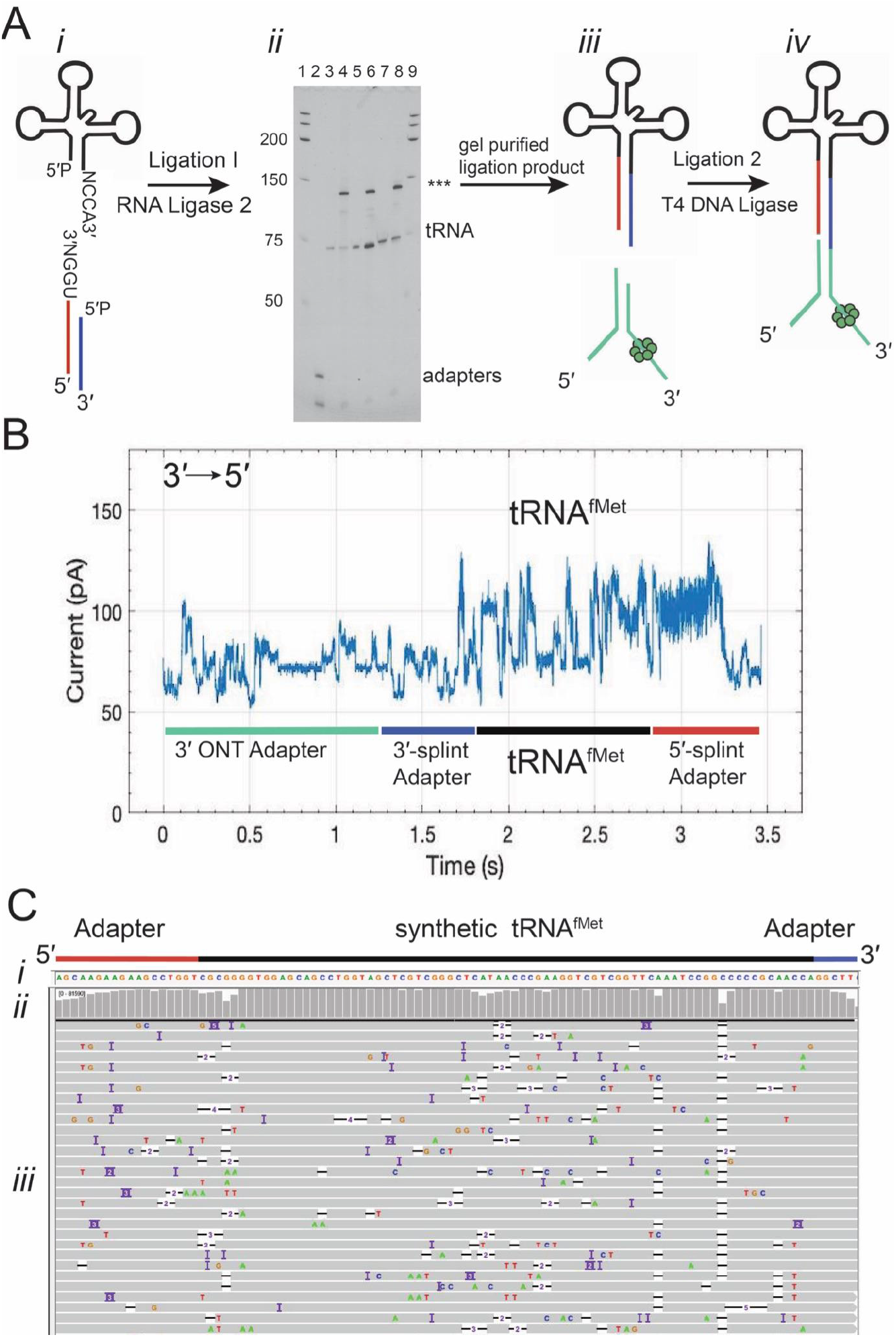
Overview of the tRNA sequencing strategy using synthetic canonical tRNAs. A) tRNA adaptation for nanopore sequencing. From left to right: *i*. The tRNA is ligated to a double-stranded splint adapter using RNA Ligase 2. *ii*. gel purification of the ligation I product for synthetic tRNAs. The denaturing PAGE gel shows the first ligation of three synthetic tRNAs to the splint adapter. The lanes are as follows 1: RNA ladder, 2: Splint adapter, 3: synthetic tRNA ^fMet^, 4: synthetic tRNA ^fMet^ ligation reaction, 5: synthetic tRNA^Lys^, 6: synthetic tRNA ^Lys^ ligation reaction, 7: synthetic tRNA^Phe^, 8: synthetic tRNA^Phe^ ligation reaction, 9: RNA ladder. The full length product (***) is excised and purified. *iii.* The purified product is ligated to the ONT sequencing adapters using T4 DNA Ligase. *iv.* Adapted, Nanopore sequencing-ready tRNA product. In the line drawing the adapters and tRNA are not to scale. B) An example ionic current trace of adapter-ligated synthetic tRNA^fMet^. Regions of the trace are indicated with colored bars corresponding to structures in A: The 3′ strand of the ONT RMX adapter (teal); the 3′ strand of the splint adapter (blue); the tRNA (black); the 5′ strand of the splint adapter (red). C) Primary alignments of synthetic tRNA^fMet^ to the reference sequence visualized using IGV. The reference sequence and its components are labeled as (*i*). The coverage at each position (coverage plot) is indicated by grey columns in (*ii*). Beneath the coverage plot is a diagram of a randomly downsized sample of the aligned reads (*iii*). Grey rows denote continuous alignment and agreement with the reference nucleotide. Within each read, positions that do not match the reference (U(T) = red, A = green, C = blue, G = gold) are shown. White spaces bisected with a black bar within an aligned read indicates a deletion. Insertions are indicated in purple. The rows of aligned reads are presented as they were displayed on IGV^17^.

The four different oligonucleotides that ligate to the 5′ end of the tRNA were identical except for the terminal base (Fig 1A *i*, red strand). Each is composed of 6 DNA followed by 18 RNA nucleotides. The 24 nt sequences of these oligonucleotides are shown below:

5′P-CCTAAGrArGrCrArArGrArArGrArArGrCrCrUrGrGrA-3′ (UCCA complement)
5′P-CCTAAGrArGrCrArArGrArArGrArArGrCrCrUrGrGrU-3′ (ACCA complement)
5′P-CCTAAGrArGrCrArArGrArArGrArArGrCrCrUrGrGrC-3′ (GCCA complement)
5′P-CCTAAGrArGrCrArArGrArArGrArArGrCrCrUrGrGrG-3′ (CCCA complement)

### Annealing splint adapters

10 μM stocks of the double-stranded splint adapters were prepared in 10mM Tris (pH 8.0), 50 mM NaCl,1 mM EDTA by adding 100 pmol of each strand in a total volume of 10 μl. The solution was heated to 75°C for 15 sec and slowly cooled to 25°C to hybridize the adapter strands.

### Library Preparation

tRNA libraries were prepared using the SQK-RNA002 (Oxford Nanopore Technologies) kit as described below.

### First Ligation: tRNA to splint adapter

In the first ligation reaction, the splint adapter to tRNA molar ratio was 1:1.25. The reaction was carried out at room temperature in a DNA LoBind tube for 45 min. Its constituents were 1X RNA Ligase 2 buffer (NEB) supplemented with 5% PEG 8000, 2 mM ATP, 6.25 mM DTT, 6.25 mM MgCl_2_ and 0.5 units/μl T4 RNA ligase 2 (10,000 units/mL). For single tRNA isotype libraries (for tRNA^fMet^, tRNA^Lys^ or tRNA^Phe^), 16 pmol of splint adapter and 20 pmol of tRNA (~500 ng), in a total reaction volume of 20 μl, was used. The tRNAs for these libraries have ACCA 3′ termini. The splint adapter used was the form with a UGGU overhang. Total tRNA reactions using all four splint adapters were performed using 32 pmol of adapter (8 pmol of each of the 4 adapters) and 40 pmol (~1 μg) of total tRNA in a reaction volume of 40 μl. For total tRNA runs using only one of the four adapters, 16 pmol of adapter and 20 pmol (~500 ng) of total tRNA were used in a reaction volume of 20 μl.

### Gel Purification of Ligation I product

Gel excision and purification is recommended, but optional for this procedure. Unligated and partially ligated tRNA carried forward to subsequent reactions may decrease the throughput and coverage. If gel purification is not done, we recommend checking a small amount (~1 μl) of the ligation reaction on a gel to validate the presence of full length product. Then bring the remaining ligation reaction up to 40 μl with NF H_2_0, clean up with 1.8X RNAclean AMPure XP beads (Novagen) and elute in a final volume of 11 μl NF H_2_0. Next, follow the procedures starting at the section “Second Ligation: splint-ligated tRNA and RMX adapter” below.

### PAGE Gel separation and excision of the tRNA/splint ligation product

The ligation reaction was diluted to 1X with 2X RNA loading dye (NEB). Standards (low range ssRNA ladder, NEB) and 10 pmol of unligated tRNA sample were prepared in an equivalent buffer to the first ligation reaction (1X RNA Ligase 2 buffer (NEB), 5% PEG 8000, 2mM ATP, 6.25 mM DTT, 6.25 mM MgCl_2_) and diluted with an equal volume of 2X RNA loading dye.

The size standard, unligated tRNA and ligation reaction samples were run on a denaturing 7M urea/TBE PAGE gel (8%) in 1X TBE buffer for ~1.5 hr at 28 Watts. The gel was post-stained in the dark, in a 1X TBE solution containing 2X SybrGold™ (Life Technologies) for 20 min. The gel was transferred to a piece of saran wrap, and using UV shadowing, the gel region corresponding to the fully ligated product (~130 nt for tRNA^fMet^, tRNA^Lys^ and tRNA^Phe^ or from ~120-170 nt for total tRNA) was excised.

### Gel purification of tRNA/splint ligation product by electroelution

The excised gel fragment was electroeluted in 1X TAE buffer using 3.5 kDa MWCO D-Tube dialyzers (Novagen) according to the manufacturer′s protocol with minor modifications. For the ethanol precipitation step, the solution was precipitated overnight at −20°C with 0.1X Sodium Acetate (pH 5.2), 1 μl glycogen (20 mg/ml, RNA grade) and 2.5-3X ethanol. Following centrifugation at 4°C for 30 min at 12,000 g, the solution was removed. 200 μl of 80% ethanol was added to wash each pellet. After centrifugation for 15 min, the ethanol was removed and the pellets were air dried briefly. The pellets were resuspended and pooled using NF H_2_0. For single isotype tRNA libraries or total tRNA libraries where one of the four adapters was used, a total of 12 μl of NF H_2_0 was used to resuspend the pellets. For total tRNA libraries where all four adapters were ligated, the resuspension volume was 24 μL of NF H_2_0. The concentration of the sample at this point may be quantified by Nanodrop or the Qubit fluorometer RNA HS assay. The amount of material recovered and carried forward to the second ligation varied between ~60-200 ng for single isotype tRNA libraries and total tRNA libraries using a single version of double stranded splint adapter. Approximately 500 ng was recovered in total tRNA libraries ligated to all 4 adapter versions. For this library, the amount of input tRNA (1 μg) was twice as much as other libraries (0.5 μg). On the order of 25% of the material by weight of the input tRNA is recovered following purification of the full length product, but this can vary substantially.

### Second Ligation: splint-ligated tRNA and RMX adapter

For single isotype tRNA libraries or total tRNA libraries where one version of the splint adapters was used, the second ligation reaction was composed of: 11 μL of the gel purified splint ligation product, 5 μL of 5X Quick Ligation Reaction buffer (NEB: B6058S), 6 μL of the RMX adapter and 3 μL of T4 DNA ligase (2,000,000 units/mL). RMX adapter is included in ONT′s RNA sequencing kits.

For total tRNA libraries previously ligated to all four types of double-stranded splint adapters, the second ligation reaction included 23 μL of the purified splint ligation products, 8 μL of 5X Quick Ligation Reaction buffer (NEB: B6058S), 6 μL of the RMX adapter and 3 μL of T4 DNA ligase (2,000,000 units/mL).

Ligation reactions were carried out at room temperature for 30 min. A 1.5X volume of Ampure RNAClean XP beads (Beckman-Coulter) was then added and mixed into the reaction by pipetting up and down. The tube was incubated for 15 min at room temperature with occasional light tapping, pelleted on the magnet, and the supernatant was removed. Two 150 μl washes with WSB (Wash buffer in the ONT kit) were conducted, during which, the pellet was vigorously resuspended by flicking, returned to the magnet to pellet, and the wash solution was removed. Following the second wash, the pellet was resuspended in 12.5 μL elution buffer (EB), and incubated for 20 min at room temperature with light tapping. Following pelleting of the beads on the magnet, the eluate was recovered to a fresh tube.

### Flow cell Quality Control (QC), priming the flow cell and loading the sample on the minION

The ONT SQK-RNA002 protocol was followed for minION flow cell (FLO-MIN-106) QC, priming, preparation of the sample in RNA running buffer (RRB) and for loading the library onto the flow cell.

### RNA handling practices

Care was made to avoid introducing RNAses into the samples or into stock solutions by wearing gloves at all times, using RNAse free filter tips and nuclease free water (NF H_2_O). Pipettes, benches and equipment were cleaned with RNAse AWAY™.

### MinION running parameters

Sequencing runs were done with live base-calling off. The experiments were set for the standard 48 hr period, but were typically run for less than 24 hrs due to a deterioration in functional channels over time seen using our samples. For sequencing runs where the nanopores in the flow cell became clogged (indicated by reduced functional pores on the MinKnow GUI), the experiment was restarted up to five times.

### Bioinformatics

#### Total tRNA reference curation

The reference used for the total tRNA experiments was designed to encompass all isoacceptor families (total tRNA reference)^14^. This included the grouping of tRNAs differing in their anticodons for all 20 amino acids, tRNA^Sec^ and the initiator tRNA (tRNA^fMet^). Composed of 42 tRNAs, it was generated from 38 modomics sequences^3^ and 4 sequences from gtRNAdb^14^. References contained the ribonucleotide portions of the splint adapter for all analyses except for the error profile assessment which contained only the tRNA sequences (Table 2). The references are shown in Supplemental Figure 1.

#### Base calling and alignment

Base calling was done with Guppy v3.0.3 using the flipflop model. The resulting reads FASTA file was then processed to convert all Us to Ts (further analysis software will not work without this step). Sequence alignment was accomplished using BWA-MEM v0.7.17-r1188 (parameter “-W 13 -k 6 -x ont2d”)^15^. The SAM files were filtered for primary alignments and for a mapping quality of > 0 (removing nonspecific alignments) using Samtools v1.6^16^, and visualized using the Broad Institute′s Integrated Genome Viewer (IGV) v2.4.14^17^. The error model statistics were calculated using marginStats v0.1^18^ (Table 2).

#### Alignment statistics

We determined the quality of our alignments using the marginAlign subprogram maginStats^18^. This program utilizes a metric called “alignment identity” which can be defined as the percent of each read (both full and partial length) that matched the reference. The equation for calculating the alignment identity is as follows:

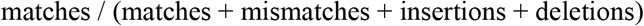

We report median alignment identity over all aligned reads (Table 2). This is distinct from “Full length Aligned Reads” (Table 1), which is the number or percentage of reads that cover the entire length of the tRNA reference sequence end-to-end.

**Table 1.** Sequencing and alignment statistics for *E. coli* tRNA^fMet^, tRNA^Lys^, and tRNA^Phe^. Alignments were generated using references which contained adapter sequences, and were filtered for a mapping quality > 0. The “Full Length” categories refer to the number or percentage of aligned reads that span the full length of the tRNA reference sequence without the adapters.

|  | Synthetic |  |  | Biological |  |  |
| --- | --- | --- | --- | --- | --- | --- |
|  | tRNA<sup>fMet</sup> | tRNA<sup>Lys</sup> | tRNA<sup>Phe</sup> | tRNA<sup>fMet</sup> | tRNA<sup>Lys</sup> | tRNA<sup>Phe</sup> |
| Number of Reads | 126, 231 | 138,110 | 141, 212 | 137,553 | 50,877 | 64,746 |
| Number of Aligned Reads | 83,956 | 103,834 | 72,870 | 67,396 | 22,087 | 36,731 |
| % of Reads that Aligned | 66.5% | 74.8% | 51.6% | 49.0% | 43.4% | 56.7% |
| Number of Aligned Reads that were Full Length | 72,034 | 72,683 | 31,188 | 51,490 | 20,342 | 33,682 |
| % of Aligned Reads that were Full Length | 85.8% | 70.0% | 42.8% | 76.4% | 92.1% | 91.7% |

**Table 2.** Read alignment error profiles for *E. coli* tRNA^fMet^, tRNA^Lys^, and tRNA^Phe^. Alignments were generated using references which did not contain adapter sequences, and were filtered for a mapping quality of > 0. The first column to the left indicates the relevant error profile metrics. The following three columns contain the error profiles for synthetic *E. coli* tRNA^fMet^, tRNA^Lys^ and tRNA^Phe^. The last three columns contain the error profiles for biological *E. coli* tRNA^fMet^, tRNA^Lys^ and tRNA^Phe^.

|  | Synthetic |  |  | Biological |  |  |
| --- | --- | --- | --- | --- | --- | --- |
|  | tRNA<sup>fMet</sup> | tRNA<sup>Lys</sup> | tRNA<sup>Phe</sup> | tRNA<sup>fMet</sup> | tRNA<sup>Lys</sup> | tRNA<sup>Phe</sup> |
| Median Alignment Identity | 86.5% | 86.3% | 84.0% | 78.9% | 75.3% | 76.1% |
| Median Read Coverage | 98.3% | 98.4% | 97.3% | 96.2% | 97.2% | 95.7% |
| Median Mismatches Per Aligned Base | 5.7% | 5.8% | 7.7% | 12.9% | 15.5% | 15.2% |
| Median Deletions Per Read Base | 4.1% | 4.1% | 4.4% | 4.0% | 4.3% | 4.1% |
| Median Insertions Per Read Base | 1.4% | 1.5% | 1.8% | 2.9% | 1.7% | 3.2% |

#### Miscall analysis

Three error models were generated with marginAlign v0.1 (EM training enabled-- BWA-MEM; parameter “-W 13 -k 6 -xont2d”) from the synthetic canonical tRNA^fMet^, tRNA^Lys^ and tRNA^Phe^ alignments. These error models (HMM file) were then used as input models to generate their corresponding biological tRNA alignments (under the same marginAlign settings, see usage on: https://github.com/mitenjain/marginAlign-tRNA). Alignments for the synthetic canonical tRNAs were also generated using their own error models, as a control^8^. The alignment files were then filtered for primary alignments using Samtools v1.6^16^. Systematic miscall identification was performed for the biological and synthetic alignments using marginCaller v0.1 (parameter “--alignmentModel= file.hmm --errorModel= file.hmm > output.vcf”) with their respective trained synthetic error models. We used a posterior probability threshold of ≥ 30% which is the default for marginCaller^18^.

For tRNA^Ala1^ miscall analysis the same strategy was applied, however the RNA-based error model was generated from canonical IVT data (Supplemental Materials and Methods).

### Liquid Chromatography Tandem Mass Spectrometry confirmation of tRNA^fMet^ modifications

Liquid chromatography tandem mass spectrometry (LC/MS-MS) with selective ion monitoring was performed to determine if the expected RNA modifications were present in biological *E. coli* tRNA^fMet^. The samples were digested to ribonucleosides using a three enzyme protocol (similar to Crain,1990)^19^. All water used was HPLC grade. One to four μg of tRNA was digested with 1 unit of nuclease P1 (Sigma Aldrich) in a 10 μL solution of 10 mM NH_4_OAc at 45°C for 2 hrs. The solution was adjusted to 50 mM NH_4_HCO_3_ and 2.5 mM MgCl_2_ and treated with 0.004 units of Phosphodiesterase 1 (Thermo Fisher Scientific) in a total volume of 20 μl at 37°C for 3 hours. 0.5 μl of 10X Antarctic phosphatase buffer and 0.5 units of Antarctic phosphatase (NEB) were added, and the solution brought to 25 μl and incubated at 37°C for 1 hr. Mock digest solutions without substrate were used to prepare the standards to maintain uniform buffer conditions. Both sample and standards were brought up to 55 μl with 0.1% formic acid and purified on 3.5 MWCO/Nanosep 3K spin columns (Pall) for 10 minutes at 14,000 RCF. The flowthrough was retained for analysis. The amount of material for LC/MS-MS runs was 0.7-1.1 μg for digested tRNA^fMet^ samples, and 60 ng for each standard.

Standards included 5-methyluridine (TCI America), *B*-pseudouridine (TRC Canada), uridine (Sigma), 4-thiouridine (MP Biomedicals), 2′-O-methylcytidine (Alfa Aesar), 7-methylguanosine (Sigma).

LC-MS/MS was done at the UC Santa Cruz Mass Spectrometry facility, with an LTQ-Orbitrap Velos Pro Mass Spectrometer (ThermoFisher) in positive ion mode. The column used was a Synergi 4 μm Fusion-RP 80Å C18 column (Phenomenex). Two solvents were used: 0.1% formic acid in H_2_O (A) and 0.1% formic acid in acetonitrile (B). The solvents gradients were: time(t) = 0-15 min: 100% A, t= 15-15.1 min 60% A, t=15.1-20.1 min: 10% A, t=20.1-30 min: 100% B. The flow rate of the chromatography was 200 μl/min.

The Xcalibur software (Thermo Fisher) was used to control the LC-MS/MS and for data analysis. Selective ion monitoring was performed and the following transitions^3,20^ were evidence of the presence of a modification: pseudouridine (245 > 209, 177, 155), 5-methyluridine (259 > 127), 4-thiouridine (261 > 129), 2′-O-methylcytidine (258 > 112), 7-methylguanosine (298 > 166). For these modifications we assessed whether the retention time for the samples was comparable to that of the standards. A commercially available dihydrouridine sample was not available, so we relied solely on published base peak (247) and daughter ion values (115) to confirm its presence.

## RESULTS

The Nanopore tRNA sequencing strategy is shown in Figure 1A: (*i*) tRNA molecules are ligated to double-stranded splint adapters (5′ adapter strand, red line; 3′ adapter strand, blue line) using RNA ligase 2; (*ii*) the ligation product is run on an 8% denaturing PAGE gel and the band corresponding to the ligated product (~130 nt indicated by asterisks) is excised and purified; and (*iii*) this purified product is ligated to the ONT motor-associated sequencing adapter using T4 DNA ligase which yields the final sequence-ready product (*iv*).

We first implemented this strategy using synthetic canonical 5′-phosphorylated tRNA^fMet^. Figure 1B shows an ionic current trace associated with translocation of an adapted tRNA strand through the nanopore. At zero seconds, the open channel current is ~245 pA (not shown). Upon strand capture the current drops to approximately 60 pA. This is followed by discrete ionic current transitions corresponding to the 3′ adapter (teal and blue bars); tRNA^fMet^ (black bar); and the 5′ splint (red bar).

Next, the nanopore ionic current data were base-called using Guppy (v3.0.3). The resulting sequences were then aligned to a reference sequence using BWA-MEM^15^. This reference sequence contained the 18 ribonucleotides of the 5′ splint adapter strand, the *E. coli* tRNA^fMet^ sequence, and the six ribonucleotides of the 3′ splint adapter strand. The resulting 83,956 aligned reads were visualized using integrated genome viewer (IGV) software^17^ (Figure 1C). Grey bars in the coverage plot (Figure 1C *i*) indicate positions where 80% or more of the quality weighted reads matched the expected nucleotide at that position.

### Sequencing purified biological and synthetic canonical tRNAs

We applied this sequencing method to commercially available purified biological *E. coli* tRNA^fMet^, tRNA^Lys^, and tRNA^Phe^ and their corresponding synthetic canonical control tRNAs. Because these tRNAs have an ACCA 3′ overhang, we used a splint adapter terminating with 5′-UrGrGrU-3′ (See Materials and Methods). Biological tRNA sequence reads and synthetic control sequence reads were aligned to their respective references using BWA-MEM and filtered for primary alignments (see Materials and Methods). The sequencing statistics and error profiles for these alignments are shown in Tables 1 and 2.

The primary alignments for biological and synthetic tRNAs are shown in Figure 2. Grey columns in the IGV coverage plots (Fig 2 a *i*, b *i*, c *i*) indicate positions where 80% or more of the reads (weighted for quality) matched the reference^17^. When the accuracy fell below this threshold, the proportion of reads for each nucleotide are indicated by the colors red, blue, gold and green for U(T), C, G and A respectively. The median alignment identity for the synthetic canonical tRNAs was 84-86%, and for the biological tRNAs it was 75-79% (Table 2).

**Figure 2.**
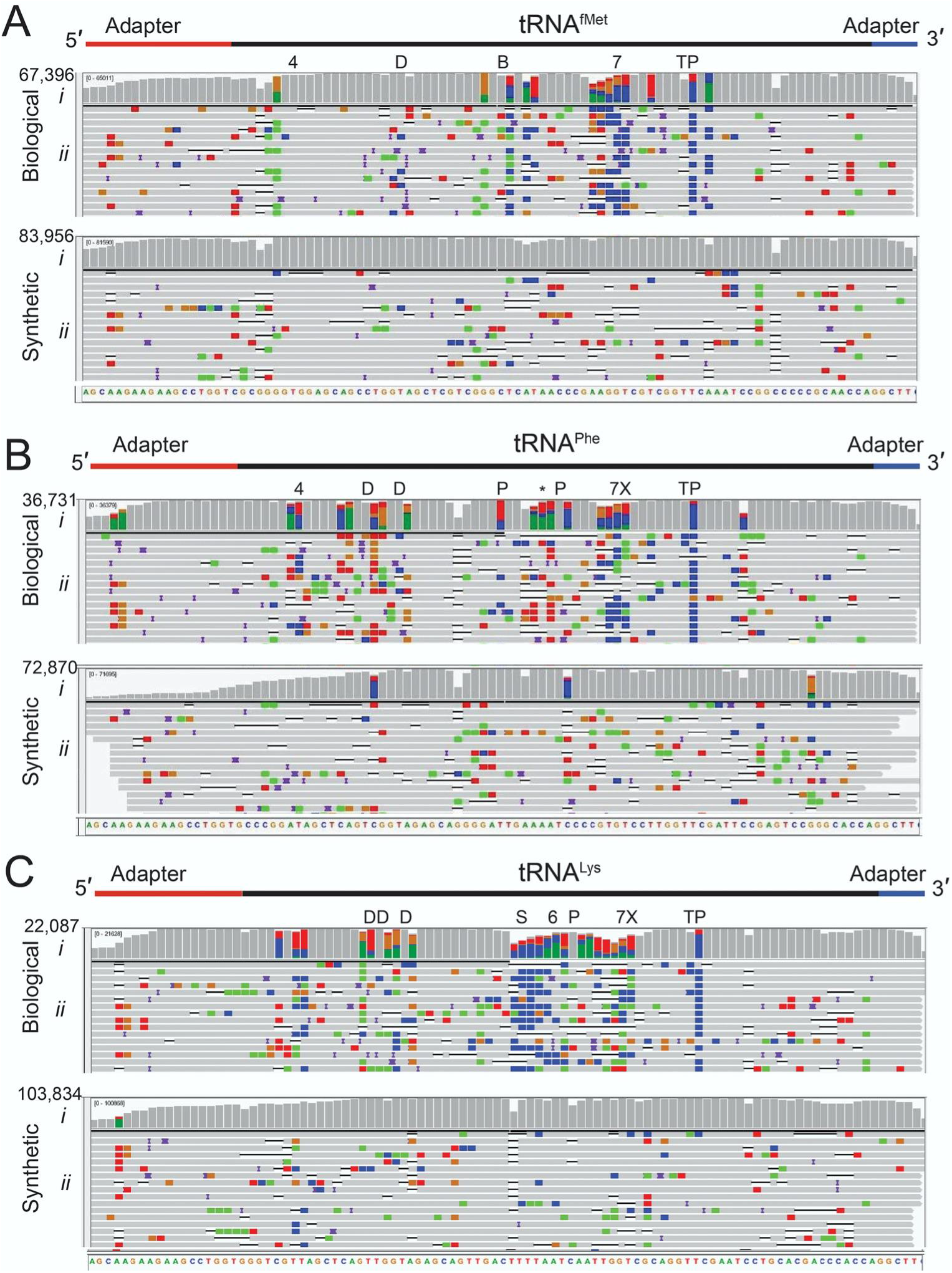
Biological and canonical tRNA strand reads aligned against reference sequences. Panel A, tRNA^fMet^; panel B tRNA^Phe^; Panel C tRNA^Lys^. In each panel (*i*) is base coverage along the reference sequence at each position (coverage plot), and (*ii*) is a randomly selected subset of individual aligned Nanopore reads. The total number of aligned reads are shown to the left of the coverage plots. The positions of expected modifications on biological tRNA^3^ are indicated above the coverage plots and are abbreviated: 4= 4-thiouridine; D= Dihydrouridine; B= 2′-O-methylcytidine; 7= 7-methylguanosine; T= 5-methyluridine; P= Pseudouridine; X= 3-(3-amino-3-carboxypropyl)uridine; * = 2-methylthio-N6-isopentenyladenosine; S= 5-methyl-aminomethyl-2-thiouridine; 6= N6-threonylcarbamoyl-adenosine. Grey columns in the coverage plots indicate positions along the reference where 80% or more of the quality weighted reads are the expected canonical nucleotide. At positions where the value is under the 80% threshold, the proportion of each nucleotide call is shown in color where U(T) = red, A = green, C = blue, and G = gold. Similarly, the rows of individual aligned reads (A-C, *ii*) are grey at positions matching the reference and colored (using the previously mentioned convention) at positions with mismatches. The black horizontal bars in the aligned reads indicate a deletion, and purple bars indicate an insertion. The rows of aligned reads are presented as they were displayed on IGV^17^.

For the biological tRNAs, the positions of known modifications^3^ are indicated above the biological tRNA coverage plots (Fig 2 A-C, *i*). The Nanopore base calls that did not match the biological tRNA references usually occured at or near the positions of known modifications.

Liquid chromatography tandem mass spectrometry (LC/MS-MS) was conducted to test for the presence of the expected modifications in biological tRNA^fMet^. Comparison of the digested nucleoside products of the tRNA to commercially available standards verified the presence of 2′-O-methylcytidine, 7-methylguanosine, 5-methyluridine and pseudouridine, but did not verify 4-thiouridine (Supplemental Table 1). The presence of dihydrouridine could not be tested using this strategy because a commercial standard is not available. However, its presence was verified based on expected parent and daughter ions.

### Systematic miscalls identified by marginCaller occurred at or near modified nucleotides

Based on the frequency of miscalls for biological tRNAs (Table 2), and their distribution in sequence alignments (Figure 2), we reasoned that some miscalls were caused by base modifications. As a test, we performed variant calling on control and biological tRNA alignments using marginCaller (Figure 3). Among miscalls, those that passed the marginCaller default posterior probability threshold ≥ 30% ^18^ were considered to be systematic miscalls.

**Figure 3.**
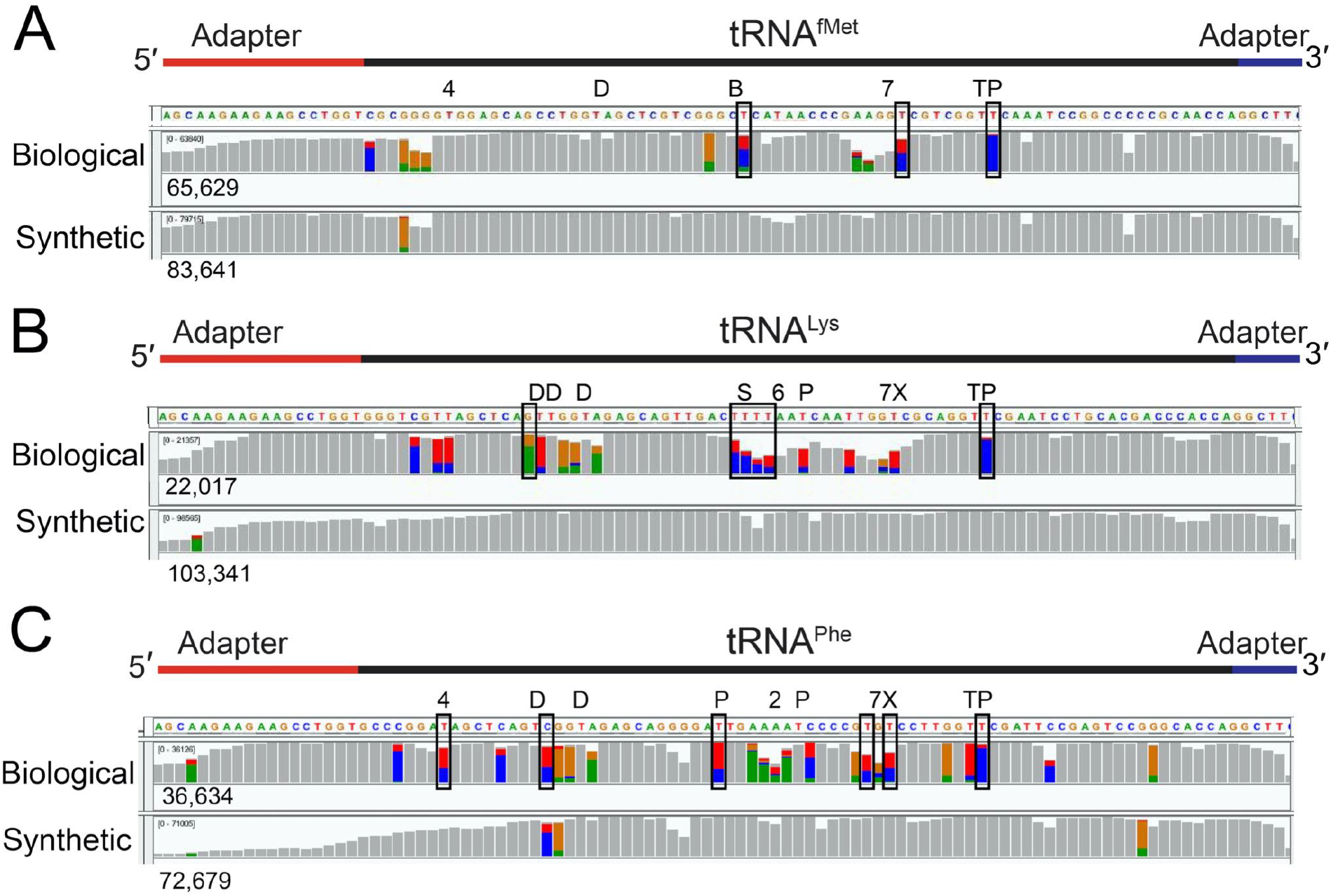
Systematic base miscalls in purified biological and canonical tRNA^fMet^, tRNA^Lys^ and tRNA^Phe^. The coverage plots (A-C) for biological and canonical synthetic tRNAs were generated from alignments using marginAlign. The number of aligned reads for each tRNA is shown under each coverage plot. Boxes surround positions of systematic miscalls (posterior probability > 30%). No systematic miscalls were identified in the synthetic canonical tRNAs. Colored bars are at positions where less that 80% of the quality weighted alignments match the reference. At these positions, the proportion of individual bases called are shown in color (U(T) =red, A = green, C = blue, and G = gold). The known modifications for the biological tRNAs^3^ are indicated above the coverage plots. Modified nucleotides are indicated above the reference sequence with abbreviations (4= 4-thiouridine, D= Dihydrouridine, B= 2′-O-methylcytidine, 7= 7-methylguanosine, T= 5-methyluridine, P = Pseudouridine, X= 3-(3-amino-3-carboxy-propyl)uridine, * = 2-methylthio-N6-isopentenyladenosine, S= 5-methylaminomethyl-2-thiouridine, 6= N6-threonylcarbamoyl-adenosine).

This method identified systematic miscalls at three positions in biological tRNA^fMet^, six positions in biological tRNA^Lys^, and six positions in biological tRNA^Phe^ (Supplemental Table 2). None of these were associated with base variants among tRNA gene copies^14^. All of these occurred within three nucleotides of a known modification. The highest posterior probability miscall in all tRNAs corresponded to pseudouridine in the T loop (Supplemental Table 2). However, not all known modification positions were detected at the default posterior probability threshold. No systematic miscalls were identified in the synthetic controls.

### Sequencing total *E. coli* tRNA

As proof of concept, we next used the Nanopore method to sequence total tRNA from *E. coli* strain MRE600. This sample included tRNAs with 4 types of 3′ NCCA overhangs (ACCA, UCCA, CCCA, and GCCA). For this reason, a combination of four double-stranded splint adapters were used for the first Nanopore sequencing ligation. This ligation product was run on a denaturing PAGE gel (Figure 4) and full-length adapted tRNA strands were excised and extracted (Figure 4, lane 4). The purified ligation I products were then ligated to the ONT sequencing adapter, sequenced on the MinION, and basecalled. The 250,542 resulting reads were then aligned to a reference set composed of the 42 *E. coli* tRNA isoacceptors^14^.

**Figure 4.**
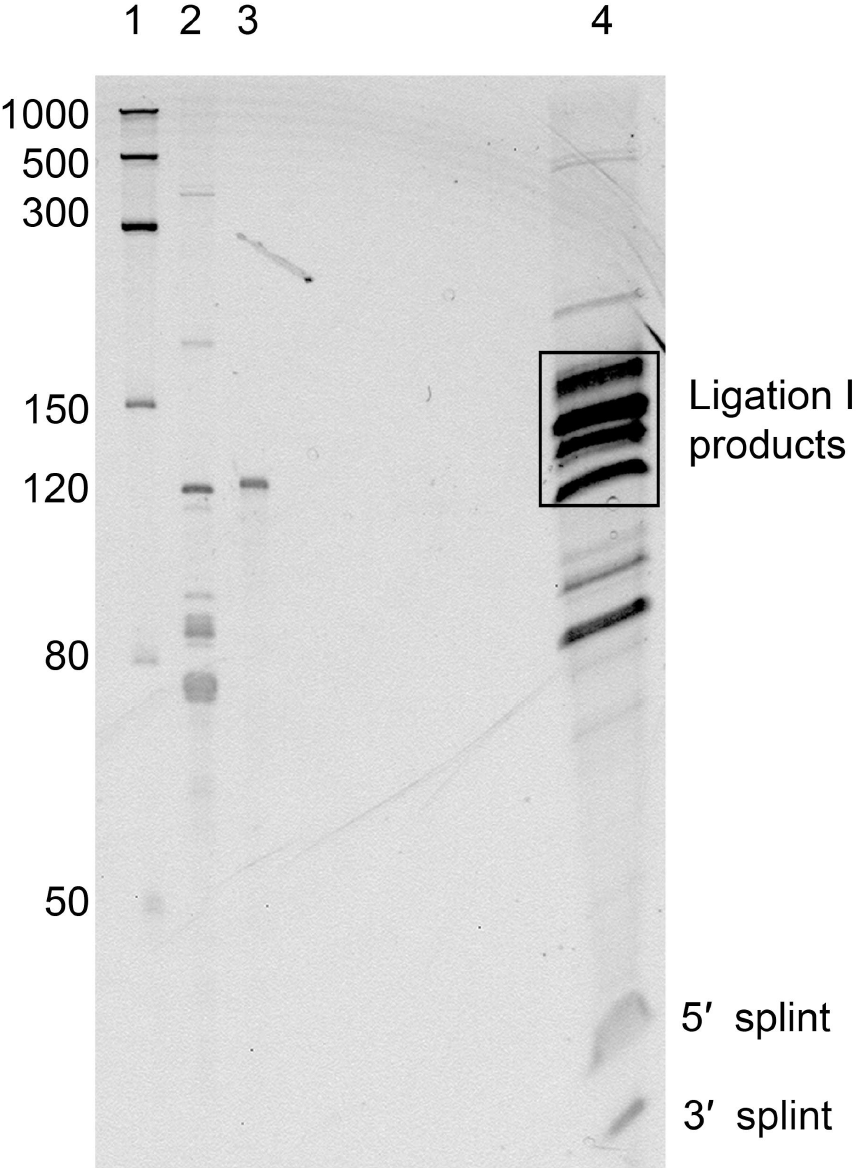
PAGE gel of total tRNA ligated to splint adapters. Lane 1, ssRNA ladder with sizes in nucleotides indicated to the left; Lane 2, *E. coli* MRE600 total tRNA; Lane 3, A 120 nt IVT 5S rRNA used as a size marker; Lane 4, the products of the ligation reaction of total *E. coli* tRNA and the 4 types of splint adapters. The two bands under 50 nt are the 30 nt and 24 nt strands of the splint adapters that did not ligate to the tRNA. Successful ligation of the double stranded splint will add 54 nt to the tRNA. As tRNA range from 75-93 nt, the expected ligation products are 129-140 nt. A block of gel encompassing fragments of approximately 110-180 nt, indicated on the gel as a black rectangle was excised, purified and carried forward for the library preparation.

Each reference tRNA sequence was appended with the RNA portion of the splint adapters to maximize recovery of aligned reads. 74,685 primary tRNA alignments were recovered of which 73,161 had a mapping quality > 0 (i.e. reads aligning uniquely to one of the sequences in the reference set^16^). Alignments were observed for all 42 isoacceptors (Table 3).

**Table 3.**
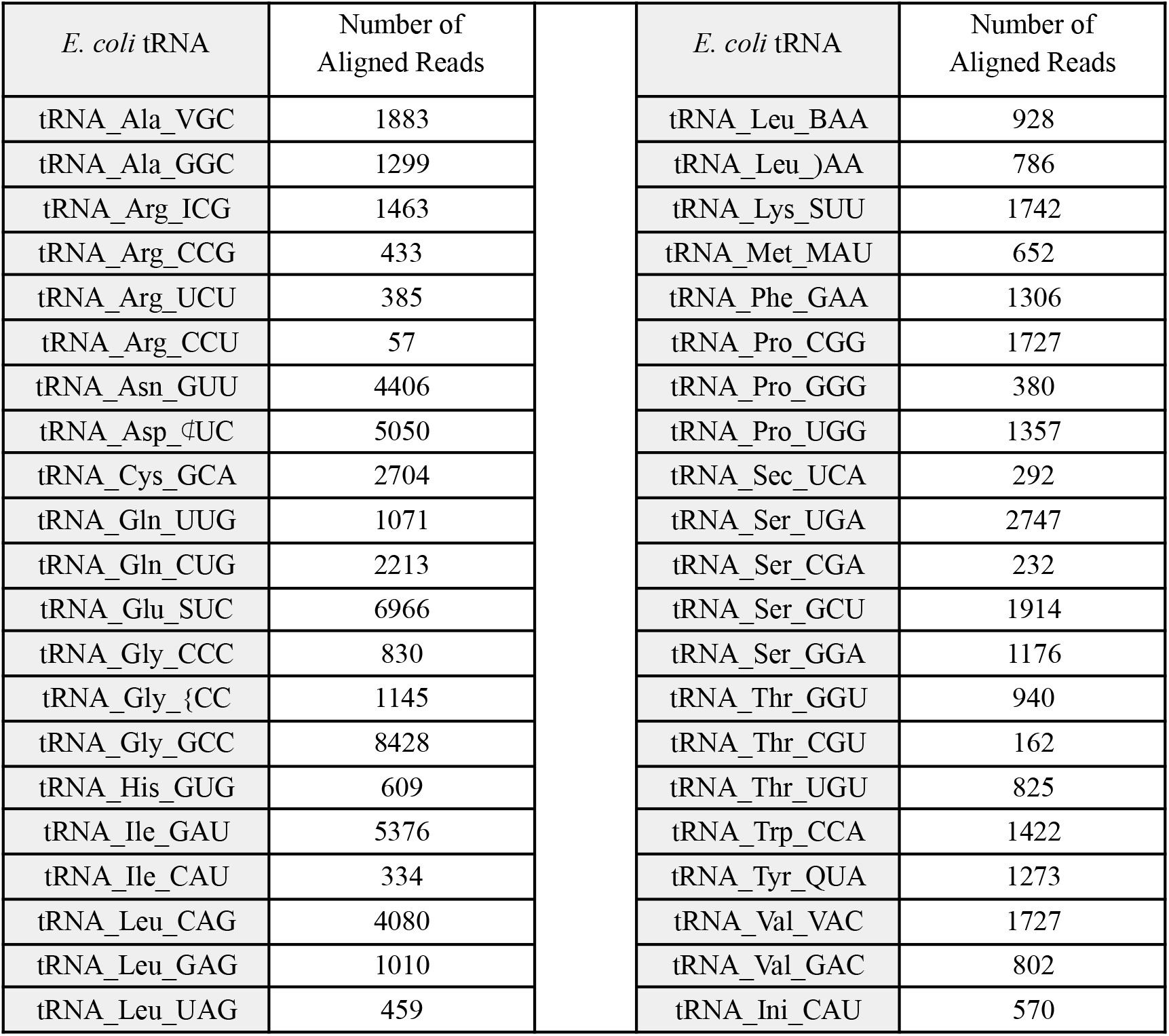
Total *E. coli* tRNA aligned reads. Values are numbers of reads aligned to each of the 42 tRNA isoacceptors with a mapping quality score > 0. The anticodon sequence is indicated for each tRNA isoacceptor. Non-canonical letters in the anticodons represent modified nucleotides^3^. Of these, 73,161 had a MAPQ > 0. These results were from total biological tRNA sequencing using all four splint adapter versions. tRNA^fMet^ (initiator tRNA) is shown as “tRNA_Ini” below.

Among the 42 *E. coli* tRNA isoacceptors, 24 end in ACCA-3′, 13 end in GCCA-3′, 4 end in UCCA-3′, and 1 ends in CCCA-3′ ^3^. We reasoned that separate tRNA capture reactions, each with a specific complimentary 4-mer overhang, would enrich for the corresponding tRNA isoacceptors. When we executed these four separate Nanopore experiments (Supplemental Table 3), we observed only a modest enrichment for tRNA with the targeted 3′ end (enrichment range 2.8%-to-33.9%; Supplemental Fig. 4). Surprisingly, reads for all 42 isoacceptor types were recovered from each experiment.

We compared our tRNA isoacceptor abundances (based on relative read numbers) to those determined by RNA fingerprinting^21^. The correlation between our results and the RNA fingerprinting was moderate (R^2^ value of 0.47, P < 0.0033 (Supplemental Figure 3)). These results were comparable to the correlation (R^2^ = 0.5 and P < 0.0001) between RNA-seq isoacceptor abundances^22^ and the same RNA fingerprinting study^21^.

We examined reads from total *E. coli* tRNA sequencing (using all adapters) for the presence of systematic miscalls using the strategy described previously (see Materials and Methods and Supplemental Table 2). As anticipated, for tRNA^fMet^, tRNA^Lys^, tRNA^Phe^ and tRNA^Ala1^ all systematic miscalls on the tRNA occurred at or adjacent to positions of known modifications^3^ (Supplemental Table 4).

### Nanopore Detection of off-target biological tRNAs

We aligned Nanopore reads for each of the purified biological tRNA samples against the total *E. coli* tRNA sequence reference set. Surprisingly, a fraction of the reads aligned to tRNAs other than the expected reference (4.6%, 7%, and 8.2% off-target reads for tRNA^fMet^, tRNA^Lys^ and tRNA^Phe^ respectively, Supplemental Table 5). This suggests that tRNA contaminants were carried over during purification of the commercial samples.

## DISCUSSION

In this paper, we used Nanopore technology to demonstrate: (i) full-length direct tRNA sequencing using custom ligated adapters; (ii) systematic miscalls at or near known modified nucleotide positions in biological tRNA alignments; and (iii) detection of all isoacceptors in total *E. coli* tRNA.

We sequenced three purified tRNAs and their matching synthetic controls. For the biological tRNAs, 76.4-92.1% of the aligned reads covered the entire tRNA. The percentage of full length synthetic tRNA reads was lower (42.8-85.8%, Table 1). The reduced coverage seen at the 5 prime end was most likely due to inefficient splint adapter ligation (Figure 3B-C). This could arise from incomplete 5′-phosphorylation of the tRNA.

Previous studies showed that Nanopore base call errors increase proximal to modified ribonucleotides^8,10,11,23^. Our data confirm this. In Table 2, median alignment identities for three biological tRNAs ranged from 75.3 to 78.9%, compared to 84 to 86.5% for their canonical controls. Furthermore, the alignment identities for the biological tRNAs were lower than for MinION-based mRNA sequencing (median 86%)^11^. This is expected because, on average, 10% of tRNA nucleotides are modified in gram negative bacteria^24^.

When we sequenced commercially purified biological tRNA samples, 49.0% of tRNA^fMet^, 43.4% of tRNA^Lys^, and 56.7% of tRNA^Phe^ reads aligned to their reference sequences. This was lower than anticipated and should improve with an updated ONT RNA basecaller that is better trained for short RNA stands. Of note, a measurable proportion of these off-target reads aligned to a different *E. coli* tRNA reference than predicted by the sample label (4.6%, 7%, and 8.2% for tRNA^fMet^, tRNA^Lys^, and tRNA^Phe^ respectively (Supplemental Table 5)). The median alignment identities for these putative tRNA impurities to alternate tRNA references ranged from 75.8% to 80.4% consistent with gene-specific tRNA alignments summarized in Table 2. We conclude that Nanopore direct tRNA sequencing could provide a fast and simple assay for tRNA purity in reference samples. This is consistent with prior work that detected 10 attomoles of an *E. coli* 16S rRNA against a background of human RNA^8^. Confirmation of this strategy will require validation using LC-MS/MS.

For nanopore sequencing of a specific tRNA in a mixture we recommend using its complementary NCCA adapter. This affords a modest, but measurable, enrichment of that target tRNA (Supplemental Figure 4). Logically, for Nanopore sequencing of all *E. coli* tRNAs in a mixture, we recommend using all four NCCA adapters. In our hands, this permitted recovery of all 42 reference isoacceptors. When we compared the relative percentages of each isoacceptor in Nanopore data to RNA fingerprinting data^21^, we found a moderate positive correlation (R^2^ = 0.47, P < 0.0033)(Supplemental Fig. 3). Although ACCA terminating tRNAs are the most abundant, comprising ~60% of *E. coli* tRNA^21^, we used an equimolar amount of each NCCA adapter. A high proportion of ACCA-terminating tRNAs (blue circles below trendline in Supplemental Fig. 3) were underrepresented in the Nanopore data relative to the RNA fingerprinting study. This suggests a limiting concentration of ACCA specific adapter in the Nanopore experiment.

We note that *E. coli* tRNA^His^ has an extra 5′ G nucleotide that base pairs in the acceptor stem resulting in a three nucleotide 3′ overhang rather than a typical four nucleotide overhang. This could account for the relatively low sequencing throughput for tRNA^His^ (Table 3), that could be resolved using a custom complementary adapter.

Every systematic miscall in our biological tRNA sequence alignments occurred within three nucleotides of a known modified position (Figure 3, Supplemental Table 2, Supplemental Table 4). The pseudouridine of the T loop (position 55) was consistently miscalled with the highest posterior probability in the purified and total biological tRNA samples that we analyzed (Supplemental Table 2, Supplemental Table 4). This tRNA modification is the most conserved modification across phyla^24^. In *E. coli*, all tRNAs have this modification at this position^3^. The possible contribution of the neighboring 5-methyluridine (position 54) to the systematic miscall remains to be explored.

Systematic miscalls were not identified at or near some known modified positions (Figure 3). Factors that could contribute to this include a modification’s chemical structure, neighboring nucleotide context, abundance at a given position, and the stringency used in defining a position as a systematic miscall. Miscall analysis is currently limited based on the constraints of the basecaller and the quality of the alignments^12,26,27^.

More accurate identification of tRNA modifications will require machine learning of associated ionic current signatures, as has been implemented for DNA methylation identification^26,28,29^. This training must include ionic current datasets for synthetic canonical tRNA, synthetic tRNA bearing known modifications at defined positions in isolation, and highly purified biological tRNA isolates. Model validation will in turn require orthogonal confirmation and mapping of modifications using LC/MS-MS. Improved nanopore base calling accuracy (which for mRNA has remained at ~87% identity for several years) will be essential.

We have presented a protocol for end-to-end sequencing of individual biological tRNA molecules using nanopore technology. We used *E. coli* tRNA because that population is well characterized. Preliminary data from our laboratory suggest that the technique will apply to tRNAs from other species. With further optimization, it could provide insights into tRNA-associated human diseases^27,30–33^.

## Acknowledgements

Andrew Smith conceived and implemented the original tRNA ligation strategy that was modified in this study. Kate Lieberman and Harry Noller provided purified tRNA^fMet^, tRNA^Lys^, and tRNA ^Phe^ samples. Li Zhang at the UCSC Mass Spectrometry Facility provided expert help and advice for detecting tRNA modifications using LC/MS-MS. Sanaiya Islam provided technical assistance. This work was supported by NIH Grant HG010053 (MA) and Oxford Nanopore Technologies Grant SC20130149 (MA).

## Declaration of interest statement

M.A. holds options in Oxford Nanopore Technologies (ONT). M.A. is a paid consultant to ONT. M.A., M.J. received reimbursement for travel, accommodation and conference fees to speak at events organised by ONT. M.A. is an inventor on 11 UC patents licensed to ONT (6,267,872, 6,465,193, 6,746,594, 6,936,433, 7,060,50, 8,500,982, 8,679,747, 9,481,908, 9,797,013, 10,059,988, and 10,081,835). M.A. received research funding from ONT.

## Data availability statement.

Ionic current and base called data will be made available upon request. They will be published in their entirety with a peer reviewed publication.

