## Supplemental Information for "Direct Nanopore Sequencing of Individual Full Length tRNA Strands"

**SUPPLEMENTAL INFORMATION**
**Supplemental Tables**
**Supplemental Figures**
**Supplemental Materials and Methods**

### SUPPLEMENTAL TABLES

**Supplementary Table 1. Summary of modifications in tRNA $^{fMet}$ verified by LC/MS-MS.**
The table shows each reported *E. coli* tRNA $^{fMet}$ modifications[1], the retention time of the standard for that modification, and for the parent ion identified in our sample, and the SRM transition monitored (positive ion mode). The presence all reported modifications with the exception of 4-Thiouridine were verified.

| Modification | Retention time standard | Retention time sample | Positive ion mode SRM Transition monitored | Verified in sample? |
|---|---|---|---|---|
| 4-Thiouridine | 9.0 min | not found | 261>129 | No |
| Dihydrouridine | N/A | 3.72 min | 247>115 | Yes |
| 2′-O-methylcytidine | 4.72 min | 4.68 min | 258>112 | Yes |
| 7-methylguanine | 4.81 min | 4.79 min | 298>166 | Yes |
| 5′methyluridine | 8.19 min | 8.17 min | 258>127 | Yes |
| Pseudouridine | 3.3 min | 3.3 min | 245>209,177,155 | Yes |

(1) Boccaletto, P. *et al.* MODOMICS: A database of RNA modification pathways. 2017 update. *Nucleic Acids Res.* **46**, D303–D307 (2018)

**Supplementary Table 2. Systematic miscalls in purified biological tRNA$^{fMet}$, tRNA$^{Lys}$ and tRNA$^{Phe}$.**

Systematic miscalls in purified biological tRNA$^{fMet}$, tRNA$^{Lys}$ and tRNA$^{Phe}$ based on alignments to the listed references. "Predicted SNV" refers to the predicted single nucleotide variant. In the column titled "Modifications", the parenthetical numbers refer to the position of the modification relative to the systematic miscall; negative values indicate position(s) upstream (to the 5′ end) and positive values indicate position(s) downstream (to the 3′ end). Modifications are considered proximal when they are within four nucleotides of the miscalled position. No positions in the synthetic tRNAs were identified as systematic miscalls.

| Reference | Position | Reference Nucleotide | Predicted SNV | Posterior Probability (BWA-MEM + EM) | Modifications[(1)] |
|---|---|---|---|---|---|
| tRNA_Ini_CAU | 34 | T | C | 53.4% | Proximal to known 2′-O-methylcytidine (-1) |
| tRNA_Ini_CAU | 48 | T | C | 62.9% | Proximal to known 7-methylguanosine (-1) |
| tRNA_Ini_CAU | 56 | T | C | 92.9% | Known pseudouridine and proximal to known 5-methyluridine (-1) |
| tRNA_Lys_SUU | 15 | G | A | 73.1% | Proximal to two dihydrouridines (+1, +2) |
| tRNA_Lys_SUU | 33 | T | C | 64.3% | Proximal to 5-methylaminomethyl-2-thiouridine (+1) and N6-threonylcarbamoyladenosine (+4) |
| tRNA_Lys_SUU | 34 | T | C | 76.9% | Known 5-methylaminomethyl-2-thiouridine, and proximal to N6-threonylcarbamoyladenosine (+3) |
| tRNA_Lys_SUU | 35 | T | C | 72.5% | Proximal to 5-methylaminomethyl-2-thiouridine (-1), N6-threonylcarbamoyladenosine (+2), and pseudouridine (+4) |
| tRNA_Lys_SUU | 36 | T | C | 41.2% | Proximal to 5-methylaminomethyl-2-thiouridine (-2), N6-threonylcarbamoyladenosine (+1), and pseudouridine (+3) |
| tRNA_Lys_SUU | 55 | T | C | 88.9% | Known Pseudouridine and proximal to known 5-methyluridine (-1) |
| tRNA_Phe_GAA | 8 | T | C | 45.9% | Known 4-thiouracil |
| tRNA_Phe_GAA | 17 | C | T | 43.6% | Proximal to two dihydrouridines (-1, +3) |
| tRNA_Phe_GAA | 32 | T | C | 32.8% | Known pseudouridine |
| tRNA_Phe_GAA | 45 | T | C | 47.2% | Proximal to known 7-methylguanosine (+1) and 3-(3-amino-3-carboxypropyl)uridine (+2) |
| tRNA_Phe_GAA | 47 | T | C | 53.9% | Known 3-(3-amino-3-carboxypropyl)uridine and proximal to 7-methylguanosine (-1) |
| tRNA_Phe_GAA | 55 | T | C | 87.6% | Known pseudouridine and proximal to known 5-methyluridine (-1) |

(1) Boccaletto, P. *et al.* MODOMICS: A database of RNA modification pathways. 2017 update. *Nucleic Acids Res.* **46**, D303–D307 (2018)

**Supplemental Table 3. Total tRNA aligned read counts using all adapters and single NCCA complementing adapters.** The number of reads with a MAPQ > 0 for the 42 isoacceptor tRNAs. Each tRNA isoacceptors with anticodon sequence is listed in the left hand column. Modified bases in anticodons are abbreviated with standard RNA modification notation. The all adapter column, a duplicate of data in Table 3, is included for comparison. The total number of aligned reads were 73161, 110918, 36821, 41353 and 178132, using all four adapters, and the individual ACCA, CCCA, GCCA and UCCA targeting adapters respectively.

| tRNA | All adapters | Adapter UGGU Target ACCA | Adapter GGGU Target CCCA | Adapter CGGU Target GCCA | Adapter AGGU Target UCCA |
|---|---|---|---|---|---|
| tRNA_Ala_VGC | 1883 | 6578 | 556 | 1441 | 3607 |
| tRNA_Ala_GGC | 1299 | 4814 | 285 | 755 | 2241 |
| tRNA_Arg_ICG | 1463 | 6020 | 199 | 996 | 2407 |
| tRNA_Arg_CCG | 433 | 239 | 97 | 250 | 366 |
| tRNA_Arg_UCU | 385 | 302 | 143 | 196 | 619 |
| tRNA_Arg_CCU | 57 | 200 | 14 | 58 | 125 |
| tRNA_Asn_GUU | 4406 | 2001 | 908 | 2287 | 6638 |
| tRNA_Asp_₵UC | 5050 | 2861 | 929 | 2666 | 4239 |
| tRNA_Cys_GCA | 2704 | 382 | 3034 | 724 | 7020 |
| tRNA_Gln_UUG | 1071 | 867 | 439 | 569 | 2405 |
| tRNA_Gln_CUG | 2213 | 1764 | 976 | 1260 | 4428 |
| tRNA_Glu_SUC | 6966 | 2551 | 1839 | 4599 | 7016 |
| tRNA_Gly_CCC | 830 | 75 | 1033 | 233 | 2293 |
| tRNA_Gly_{CC | 1145 | 232 | 1103 | 211 | 5316 |
| tRNA_Gly_GCC | 8428 | 1616 | 8720 | 1869 | 31971 |
| tRNA_His_GUG | 609 | 45 | 1154 | 139 | 483 |
| tRNA_Ile_GAU | 5376 | 11129 | 1340 | 2572 | 9124 |
| tRNA_Ile_CAU | 334 | 769 | 144 | 204 | 660 |
| tRNA_Leu_CAG | 4080 | 10214 | 3173 | 2346 | 16970 |
| tRNA_Leu_GAG | 1010 | 3186 | 1229 | 719 | 1441 |
| tRNA_Leu_UAG | 459 | 1118 | 499 | 292 | 854 |
| tRNA_Leu_BAA | 928 | 1785 | 757 | 482 | 3131 |
| tRNA_Leu_)AA | 786 | 2394 | 880 | 515 | 1410 |
| tRNA_Lys_SUU | 1742 | 5148 | 677 | 1009 | 4038 |
| tRNA_Met_MAU | 652 | 1969 | 203 | 316 | 1321 |
| tRNA_Phe_GAA | 1306 | 2400 | 170 | 963 | 2073 |
| tRNA_Pro_CGG | 1727 | 2570 | 341 | 1223 | 2938 |
| tRNA_Pro_GGG | 380 | 584 | 95 | 512 | 770 |
| tRNA_Pro_UGG | 1357 | 1969 | 252 | 838 | 3013 |
| tRNA_Sec_UCA | 292 | 827 | 158 | 174 | 8842 |
| tRNA_Ser_UGA | 2747 | 4588 | 1338 | 1867 | 4166 |
| tRNA_Ser_CGA | 232 | 263 | 59 | 187 | 376 |
| tRNA_Ser_GCU | 1914 | 5440 | 867 | 1592 | 17435 |
| tRNA_Ser_GGA | 1176 | 1061 | 328 | 735 | 1275 |
| tRNA_Thr_GGU | 940 | 3391 | 121 | 709 | 1555 |
| tRNA_Thr_CGU | 162 | 664 | 21 | 118 | 420 |
| tRNA_Thr_UGU | 825 | 1614 | 194 | 447 | 1626 |
| tRNA_Trp_CCA | 1422 | 1029 | 509 | 764 | 1963 |
| tRNA_Tyr_QUA | 1273 | 4423 | 1072 | 1072 | 3744 |
| tRNA_Val_VAC | 1727 | 6556 | 502 | 1272 | 3422 |
| tRNA_Val_GAC | 802 | 2966 | 139 | 538 | 1496 |
| tRNA_Ini_CAU | 570 | 2314 | 324 | 1634 | 2895 |
| Total | 73161 | 110918 | 36821 | 41353 | 178132 |

**Supplemental Table 4. Systematic miscalls in tRNA$^{fMet}$, tRNA$^{Lys}$, tRNA$^{Phe}$, and tRNA$^{Ala1}$ from total *E.coli* tRNA.** Systematic miscalls in tRNA$^{fMet}$, tRNA$^{Lys}$, tRNA$^{Phe}$, and tRNA$^{Ala1}$ reads from total tRNA, based on alignments to the listed references. "Predicted SNV" refers to the predicted single nucleotide variant. In the column titled "Modifications", the parenthetical numbers refer to the position of the modification relative to the systematic miscall; negative values indicate position(s) upstream (to the 5′ end) and positive values indicate position(s) downstream (to the 3′ end). Modifications are considered proximal when they are within four nucleotides of the miscalled position. No positions in the synthetic tRNAs were identified as systematic miscalls.

| Reference | Position | Reference Nucleotide | Predicted SNV | Posterior Probability (BWA-MEM + EM) | Modifications[1] |
|---|---|---|---|---|---|
| tRNA_Ala_VGC | 47 | T | C | 44.1% | Proximal to 7-methylguanosine(-1) |
| tRNA_Ala_VGC | 55 | T | C | 92.8% | Known Pseudouridine |
| tRNA_Ini_CAU | 34 | T | C | 58.3% | Proximal to known 2'-O-methylcytidine (-1) |
| tRNA_Ini_CAU | 48 | T | C | 85.1% | Proximal to known 7-methylguanosine (-1) |
| tRNA_Ini_CAU | 56 | T | C | 94.1% | Known Pseudouridine, proximal to known 5-methyluridine (-1) |
| tRNA_Lys_SUU | 15 | G | A | 76.2% | Proximal to two Dihydrouridines (+1, +2) |
| tRNA_Lys_SUU | 33 | T | C | 83.4% | Proximal to 5-methylaminomethyl-2-thiouridine (+1) and N6-threonylcarbamoyladenosine (+4) |
| tRNA_Lys_SUU | 34 | T | C | 88.7% | Known 5-methylaminomethyl-2-thiouridine and proximal to N6-threonylcarbamoyladenosine(+3) |
| tRNA_Lys_SUU | 35 | T | C | 87.4% | Proximal to 5-methylaminomethyl-2-thiouridine (-1), proximal to N6-threonylcarbamoyladenosine (+2), and proximal to Pseudouridine (+4) |
| tRNA_Lys_SUU | 36 | T | C | 62.4% | Proximal to 5-methylaminomethyl-2-thiouridine (-2), proximal to N6-methyl-adenosine (+1), and proximal to Pseudouridine (+3) |
| tRNA_Lys_SUU | 39 | T | C | 40.6% | Known pseudouridine, proximal to N6-methyl-adenosine (-2) |
| tRNA_Lys_SUU | 43 | T | C | 42.4% | Proximal to pseudouridine (-4), 7-methylguanosine (+3), and 3-(3-amino-3-carboxypropyl)uridine (+4) |
| tRNA_Lys_SUU | 47 | T | C | 45.4% | Known 3-(3-amino-3-carboxypropyl)uridine and proximal to 7-methylguanosine (-1) |
| tRNA_Lys_SUU | 55 | T | C | 91.7% | Known Pseudouridine, proximal to known 5-methyluridine (-1) |
| tRNA_Phe_GAA | 8 | T | C | 61.6% | Known 4-thiouracil |
| tRNA_Phe_GAA | 17 | C | T | 51.7% | Proximal to two dihydrouridines (-1,+3) |
| tRNA_Phe_GAA | 32 | T | C | 33.5% | Known pseudouridine |
| tRNA_Phe_GAA | 37 | A | T | 44.5% | Known 2-methylthio-N6-isopentenyladenosine, proximal to pseudouridine (+2) |
| tRNA_Phe_GAA | 45 | T | C | 73.4% | Proximal to known 7-methylguanosine (+1) and 3-(3-amino-3-carboxypropyl)uridine (+2) |
| tRNA_Phe_GAA | 47 | T | C | 76.2% | Known 3-(3-amino-3-carboxypropyl)uridine, and proximal to 7-methylguanosine (-1) |
| tRNA_Phe_GAA | 55 | T | C | 92.3% | Known Pseudouridine, and proximal to known 5-methyluridine (-1) |

(1) Boccaletto, P. *et al.* MODOMICS: A database of RNA modification pathways. 2017 update. *Nucleic Acids Res.* **46**, D303–D307 (20

**Supplemental Table 5. Off-target tRNA in biological tRNA$^{fMet}$, tRNA$^{Lys}$, and tRNA$^{Phe}$ sequencing experiments.** Reads from each individual biological tRNA experiment were aligned against a reference set containing all 42 *E. coli* tRNA isoacceptors. Reads that did not align to the expected tRNA reference were realigned to the specific tRNA$^{fMet}$, tRNA$^{Lys}$, and tRNA$^{Phe}$ reference sequences respectively. Unaligned reads from this re-examination were used for tRNA impurity count analysis (see Supplemental Figure 5).

| tRNA | tRNA$^{fMet}$ Experiment | tRNA$^{Lys}$ Experiment | tRNA$^{Phe}$ Experiment |
|---|---|---|---|
| tRNA_Ala_VGC | 2 | 142 | 103 |
| tRNA_Ala_GGC | 0 | 38 | 212 |
| tRNA_Arg_ICG | 1 | 30 | 10 |
| tRNA_Arg_CCG | 1 | 3 | 23 |
| tRNA_Arg_UCU | 1 | 7 | 24 |
| tRNA_Arg_CCU | 0 | 0 | 0 |
| tRNA_Asn_GUU | 0 | 103 | 21 |
| tRNA_Asp_¢UC | 0 | 143 | 278 |
| tRNA_Cys_GCA | 4 | 3 | 11 |
| tRNA_Gln_UUG | 0 | 56 | 83 |
| tRNA_Gln_CUG | 0 | 54 | 30 |
| tRNA_Glu_SUC | 1 | 88 | 144 |
| tRNA_Gly_CCC | 0 | 4 | 2 |
| tRNA_Gly_{CC | 0 | 2 | 2 |
| tRNA_Gly_GCC | 5 | 60 | 99 |
| tRNA_His_GUG | 1 | 5 | 14 |
| tRNA_Ile_GAU | 3964 | 864 | 868 |
| tRNA_Ile_CAU | 1404 | 9 | 12 |
| tRNA_Leu_CAG | 2 | 0 | 1 |
| tRNA_Leu_GAG | 9 | 0 | 0 |
| tRNA_Leu_UAG | 1 | 0 | 1 |
| tRNA_Leu_BAA | 2 | 2 | 1244 |
| tRNA_Leu_)AA | 1 | 0 | 1 |
| tRNA_Lys_SUU | 189 | N/A | 1138 |
| tRNA_Met_MAU | 586 | 14 | 16 |
| tRNA_Phe_GAA | 3 | 1051 | N/A |
| tRNA_Pro_CGG | 1 | 49 | 187 |
| tRNA_Pro_GGG | 1 | 11 | 14 |
| tRNA_Pro_UGG | 0 | 34 | 0 |
| tRNA_Sec_UCA | 1 | 0 | 0 |
| tRNA_Ser_UGA | 16 | 0 | 0 |
| tRNA_Ser_CGA | 35 | 0 | 1 |
| tRNA_Ser_GCU | 1 | 0 | 0 |
| tRNA_Ser_GGA | 0 | 0 | 0 |
| tRNA_Thr_GGU | 1 | 6 | 7 |
| tRNA_Thr_CGU | 0 | 4 | 11 |
| tRNA_Thr_UGU | 3 | 6 | 13 |
| tRNA_Trp_CCA | 2 | 293 | 91 |
| tRNA_Tyr_QUA | 4 | 10 | 0 |
| tRNA_Val_VAC | 1 | 359 | 27 |
| tRNA_Val_GAC | 21 | 35 | 160 |
| tRNA_Ini_CAU | N/A | 63 | 349 |
| **Total (% of all reads):** | 6264 (4.6%) | 3557 (7%) | 5287 (8.2%) |

### SUPPLEMENTAL FIGURES

**Supplemental Figure 1. tRNA references used for alignments.** Each tRNA reference is appended with the RNA portions of the 5' and 3' splint adapters (highlighted in gray).

References for individual tRNAs:

```
>tRNA_Ini_CAU_Escherichiacoli_prokaryoticcytosol
AGCAAGAAGAAGCCTGGTCGCGGGGTGGAGCAGCCTGGTAGCTCGTCGGGCTCATAACCCGAAGGTCGTCGGTTCAAATCCGGCCCCCGCAACCAGGCTTC

>tRNA_Phe_GAA_Escherichiacoli_prokaryoticcytosol
AGCAAGAAGAAGCCTGGTGCCCGGATAGCTCAGTCGGTAGAGCAGGGGATTGAAAATCCCCGTGTCCTTGGTTCGATTCCGAGTCCGGGCACCAGGCTTC

>tRNA_Lys_SUU_Escherichiacoli_prokaryoticcytosol
AGCAAGAAGAAGCCTGGTGGGTCGTTAGCTCAGTTGGTAGAGCAGTTGACTTTTAATCAATTGGTCGCAGGTTCGAATCCTGCACGACCCACCAGGCTTC
```

Total tRNA reference (42 isoacceptors each appended with the RNA portions of the splint adapters):

```
>tRNA_Ala_VGC_Escherichiacoli_prokaryoticcytosol
AGCAAGAAGAAGCCTGGTGGGGCTATAGCTCAGCTGGGAGAGCGCCTGCTTTGCACGCAGGAGGTCTGCGGTTCGATCCCGCATAGCTCCACCAGGCTTC
>tRNA_Ala_GGC_Escherichiacoli_prokaryoticcytosol
AGCAAGAAGAAGCCTGGTGGGGCTATAGCTCAGCTGGGAGAGCGCTTGCATGGCATGCAAGAGGTCAGCGGTTCGATCCCGCTTAGCTCCACCAGGCTTC
>tRNA_Arg_ICG_Escherichiacoli_prokaryoticcytosol
AGCAAGAAGAAGCCTGGTGCATCCGTAGCTCAGCTGGATAGAGTACTCGGCTACGAACCGAGCGGTCGGAGGTTCGAATCCTCCCGGATGCACCAGGCTTC
>tRNA_Arg_CCG_Escherichiacoli_prokaryoticcytosol
AGCAAGAAGAAGCCTGGCGCGCCCGTAGCTCAGCTGGATAGAGCGCTGCCCTCCGGAGGCAGAGGTCTCAGGTTCGAATCCTGTCGGGCGCGCCAGGCTTC
>tRNA_Arg_UCU_Escherichiacoli_prokaryoticcytosol
AGCAAGAAGAAGCCTGGCGCGCCCTTAGCTCAGTTGGATAGAGCAACGACCTTCTAAGTCGTGGGCCGCAGGTTCGAATCCTGCAGGGCGCGCCAGGCTTC
>tRNA_Asn_GUU_Escherichiacoli_prokaryoticcytosol
AGCAAGAAGAAGCCTGGCTCCTCTGTAGTTCAGTCGGTAGAACGGCGGACTGTTAATCCGTATGTCACTGGTTCGAGTCCAGTCAGAGGAGCCAGGCTTC
>tRNA_Asp_tUC_Escherichiacoli_prokaryoticcytosol
AGCAAGAAGAAGCCTGGCGGAGCGGTAGTTCAGTCGGTTAGAATACCTGCCTGTCACGCAGGGGGTCGCGGGTTCGAGTCCCGTCCGTTCCGCCAGGCTTC
>tRNA_Cys_GCA_Escherichiacoli_prokaryoticcytosol
AGCAAGAAGAAGCCTGGAGGCGCGTTAACAAAGCGGTTATGTAGCGGATTGCAAATCCGTCTAGTCCGGTTCGACTCCGGAACGCGCCTCCAGGCTTC
>tRNA_Gln_UUG_Escherichiacoli_prokaryoticcytosol
AGCAAGAAGAAGCCTGGCTGGGGTATCGCCAAGCGGTAAGGCACCGGTTTTTGATACCGGCATTCCCTGGTTCGAATCCAGGTACCCCAGCCAGGCTTC
>tRNA_Gln_CUG_Escherichiacoli_prokaryoticcytosol
AGCAAGAAGAAGCCTGGCTGGGGTATCGCCAAGCGGTAAGGCACCGGATTCTGATTCCGGCATTCCGAGGTTCGAATCCTCGTACCCCAGCCAGGCTTC
>tRNA_Glu_SUC_Escherichiacoli_prokaryoticcytosol
AGCAAGAAGAAGCCTGGCGTCCCCTTCGTCTAGAGGCCCAGGACACCGCCCTTTCACGGCGGTAACAGGGGTTCGAATCCCCTAGGGGACGCCAGGCTTC
>tRNA_Gly_CCC_Escherichiacoli_prokaryoticcytosol
AGCAAGAAGAAGCCTGGAGCGGGCGTAGTTCAATGGTAGAACGAGAGCTTCCCAAGCTCTATACGAGGGTTCGATTCCCTTCGCCCGCTCCAGGCTTC
```

>tRNA_Gly_lcbCC_Escherichiacoli_prokaryoticcytosol
AGCAAGAAGAAGCCTGGAGCGGGCATCGTATAATGGCTATTACCTCAGCCTTCCAAGCTGATGATGCGGGTTCGATTCCCGCTGCCCGCTCCAGGCTTC
>tRNA_Gly_GCC_Escherichiacoli_prokaryoticcytosol
AGCAAGAAGAAGCCTGGAGCGGGAATAGCTCAGTTGGTAGAGCACGACCTTGCCAAGGTCGGGGTCGCGAGTTCGAGTCTCGTTTCCCGCTCCAGGCTTC
>tRNA_His_GUG_Escherichiacoli_prokaryoticcytosol
AGCAAGAAGAAGCCTGGGGGTGGCTATAGCTCAGTTGGTAGAGCCCTGGATTGTGATTCCAGTTGTCGTGGGTTCGAATCCCATTAGCCACCCCAGGCTTC
>tRNA_Ile_GAU_Escherichiacoli_prokaryoticcytosol
AGCAAGAAGAAGCCTGGTAGGCTTGTAGCTCAGGTGGTTAGAGCGCACCCCTGATAAGGGTGAGGTCGGTGGTTCAAGTCCACTCAGGCCTACCAGGCTTC
>tRNA_Ile_CAU_Escherichiacoli_prokaryoticcytosol
AGCAAGAAGAAGCCTGGTGGCCCCTTAGCTCAGTGGTTAGAGCAGGCGACTCATAATCGCTTGGTCGCTGGTTCAAGTCCAGCAGGGGCCACCAGGCTTC
>tRNA_Leu_CAG_Escherichiacoli_prokaryoticcytosol
AGCAAGAAGAAGCCTGGTGCGAAGGTGGCGGAATTGGTAGACGCGCTAGCTTCAGGTGTTAGTGTCCTTACGGACGTGGGGGTTCAAGTCCCCCCCCTCGCACCAGGCTTC
>tRNA_Leu_GAG_Escherichiacoli_prokaryoticcytosol
AGCAAGAAGAAGCCTGGTGCCGAGGTGGTGGAATTGGTAGACACGCTACCTTGAGGTGGTAGTGCCCAATAGGGCTTACGGGTTCAAGTCCCGTCCTCGGTACCAGGCTTC
>tRNA_Leu_rpAA_Escherichiacoli_prokaryoticcytosol
AGCAAGAAGAAGCCTGGTGCCCGGATGGTGGAATCGGTAGACACAAGGGATTTAAAATCCCTCGGCGTTCGCGCTGTGCGGGTTCAAGTCCCGCTCCGGGTACCAGGCTTC
>tRNA_Leu_BAA_Escherichiacoli_prokaryoticcytosol
AGCAAGAAGAAGCCTGGTGCCGAAGTGGCGAAATCGGTAGACGCAGTTGATTCAAAATCAACCGTAGAAATACGTGCCGGTTCGAGTCCGGCCTTCGGCACCAGGCTTC
>tRNA_Lys_SUU_Escherichiacoli_prokaryoticcytosol
AGCAAGAAGAAGCCTGGTGGGTCGTTAGCTCAGTTGGTAGAGCAGTTGACTTTTAATCAATTGGTCGCAGGTTCGAATCCTGCACGACCCACCAGGCTTC
>tRNA_Met_MAU_Escherichiacoli_prokaryoticcytosol
AGCAAGAAGAAGCCTGGTGGCTACGTAGCTCAGTTGGTTAGAGCACATCACTCATAATGATGGGGTCACAGGTTCGAATCCCGTCGTAGCCACCAGGCTTC
>tRNA_Phe_GAA_Escherichiacoli_prokaryoticcytosol
AGCAAGAAGAAGCCTGGTGCCCGGATAGCTCAGTCGGTAGAGCAGGGGATTGAAAATCCCCGTGTCCTTGGTTCGATTCCGAGTCCGGGCACCAGGCTTC
>tRNA_Pro_CGG_Escherichiacoli_prokaryoticcytosol
AGCAAGAAGAAGCCTGGTCGGTGATTGGCGCAGCCTGGTAGCGCACTTCGTTCGGGACGAAGGGGTCGGAGGTTCGAATCCTCTATCACCGACCAGGCTTC
>tRNA_Sec_UCA_Escherichiacoli_prokaryoticcytosol
AGCAAGAAGAAGCCTGGCAAGATCGTCGTCTCCGGTGAGGCGGCTGGACTTCAAATCCAGTTGGGGCCGCGCGGTCCCGGGCAGGTTCGACTCCTGTGATCTTGCCAGGCTTC
>tRNA_Ser_UGA_Escherichiacoli_prokaryoticcytosol
AGCAAGAAGAAGCCTGGCGGAAGTGTGGCCGAGCGGTTGAAGGCACCGGTCTTGAAAACCGGCGACCCGAAAGGGTTCCAGAGTTCGAATCTCTGCGCTTCCGCCAGGCTTC
>tRNA_Ser_CGA_Escherichiacoli_prokaryoticcytosol
AGCAAGAAGAAGCCTGGCGGAGAGATGCCGGAGCGGCTGAACGGACCGGTCTCGAAAACCGGAGTAGGGGCAACTCTACCGGGGGTTCAAATCCCCCTCTCTCCGCCAGGCTTC
>tRNA_Ser_GCU_Escherichiacoli_prokaryoticcytosol
AGCAAGAAGAAGCCTGGCGGTGAGGTGGCCGAGAGGCTGAAGGCGCTCCCCTGCTAAGGGAGTATGCGGTCAAAAGCTGCATCCGGGGTTCGAATCCCCGCCTCACCGCCAGGCTTC
>tRNA_Ser_GGA_Escherichiacoli_prokaryoticcytosol
AGCAAGAAGAAGCCTGGCGGTGAGGTGTCCGAGTGGTTGAAGGAGCACGCCTGGAAAGTGTGTATACGGCAACGTATCGGGGGTTCGAATCCCCCCCTCACCGCCAGGCTTC
>tRNA_Thr_GGU_Escherichiacoli_prokaryoticcytosol
AGCAAGAAGAAGCCTGGTGCTGATATGGCTCAGTTGGTAGAGCGCACCCTTGGTAAGGGTGAGGTCCCCAGTTCGACTCTGGGTATCAGCACCAGGCTTC
>tRNA_Trp_CCA_Escherichiacoli_prokaryoticcytosol
AGCAAGAAGAAGCCTGGCAGGGGCGTAGTTCAATTGGTAGAGCACCGGTCTCCAAAACCGGGTGTTGGGAGTTCGAGTCTCTCCGCCCCTGCCAGGCTTC
>tRNA_Tyr_QUA_Escherichiacoli_prokaryoticcytosol
AGCAAGAAGAAGCCTGGTGGTGGGGTTCCCGAGCGGCCAAAGGGAGCAGACTGTAAATCTGCCGTCATCGACTTCGAAGGTTCGAATCCTTCCCCCACCACCAGGCTTC

>tRNA_Val_VAC_Escherichiacoli_prokaryoticcytosol
AGCAAGAAGAAGCCTGGTGGGTGATTAGCTCAGCTGGGAGAGCACCTCCCTTACAAGGAGGGGGTCGGCGGTTCGATCCCGTCATCACCCACCAGGCTTC
>tRNA_Val_GAC_Escherichiacoli_prokaryoticcytosol
AGCAAGAAGAAGCCTGGTGCGTCCGTAGCTCAGTTGGTTAGAGCACCACCTTGACATGGTGGGGGTCGGTGGTTCGAGTCCACTCGGACGCACCAGGCTTC
>tRNA_Ini_CAU_Escherichiacoli_prokaryoticcytosolll
AGCAAGAAGAAGCCTGGTCGCGGGGTGGAGCAGCCTGGTAGCTCGTCGGGCTCATAACCCGAAGGTCGTCGGTTCAAATCCGGCCCCCGCAACCAGGCTTC
>tRNA_Leu_UAG_Escherichiacoli_prokaryoticcytosol
AGCAAGAAGAAGCCTGGTGCGGGAGTGGCGAAATTGGTAGACGCACCAGATTTAGGTTCTGGCGCCGCAAGGTGTGCGAGTTCAAGTCTCGCCTCCCGCACCAGGCTTC
>tRNA_Thr_UGU_Escherichiacoli_prokaryoticcytosol
AGCAAGAAGAAGCCTGGTGCCGACTTAGCTCAGTAGGTAGAGCAACTGACTTGTAATCAGTAGGTCACCAGTTCGATTCCGGTAGTCGGCACCAGGCTTC
>tRNA_Arg_CCT_Escherichiacoli_prokaryoticcytosol
AGCAAGAAGAAGCCTGGTGTCCTCTTAGTTAAATGGAtATAACGAGCCCCTCCTAAGGGCTAAtTGCAGGTTCGATTCCTGCAGGGGACACCACCAGGCTTC
>tRNA_Pro_GGG_Escherichiacoli_prokaryoticcytosol
AGCAAGAAGAAGCCTGGTCGGCACGTAGCGCAGCCtGGTAGCGCACCGTCATGGGGTGTCGGGGGtCGGAGGTTCAAATCCTCTCGTGCCGACCACCAGGCTTC
>tRNA_Pro_TGG_Escherichiacoli_prokaryoticcytosol
AGCAAGAAGAAGCCTGGTCGGCGAGTAGCGCAGCTtGGTAGCGCAACTGGTTTGGGACCAGTGGGtCGGAGGTTCGAATCCTCTCTCGCCGACCACCAGGCTTC
>tRNA_Thr_CGT_Escherichiacoli_prokaryoticcytosol
AGCAAGAAGAAGCCTGGTGCCGATATAGCTCAGTTGGTAGAGCAGCGCATTCGTAATGCGAAGGTCGTAGGTTCGACTCCTATTATCGGCACCAGGCTTC

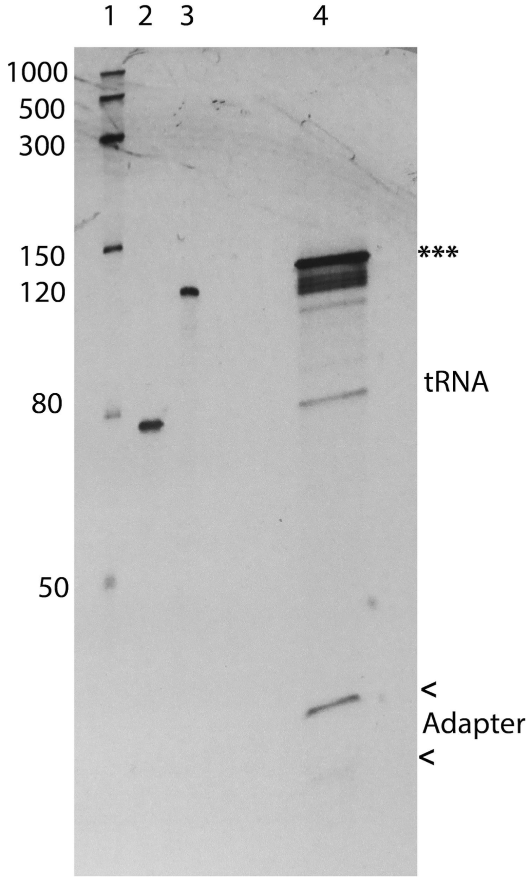


**Supplemental Figure 2**. **Biological tRNA$^{fMet}$ Ligation I Reaction.** The PAGE gel shows: Lane 1: RNA size marker, Lane 2: biological tRNA$^{fMet}$, Lane 3: 120nt IVT marker, Lane 4: The ligation reaction of tRNA$^{fMet}$ with the splint adapter. For Lane 4 it is clear that the reaction did not go to completion. Unligated adapters (indicated with carrots at 30nt and 24nt) and tRNA (76nt) are seen. The fully ligated ~130nt product, which is gel purified, is indicated with 3 asterisks. Partially ligated products between 80 and 120nt are seen. The position of unligated tRNA and unligated adapters (see carrots) are shown to the left of the gel. In some ligation reactions, products higher than 130nt may be seen. The strong band at ~130nt is excised and carried forward for the library preparation.

### tRNA Abundance: Nanopore vs RNA fingerprinting

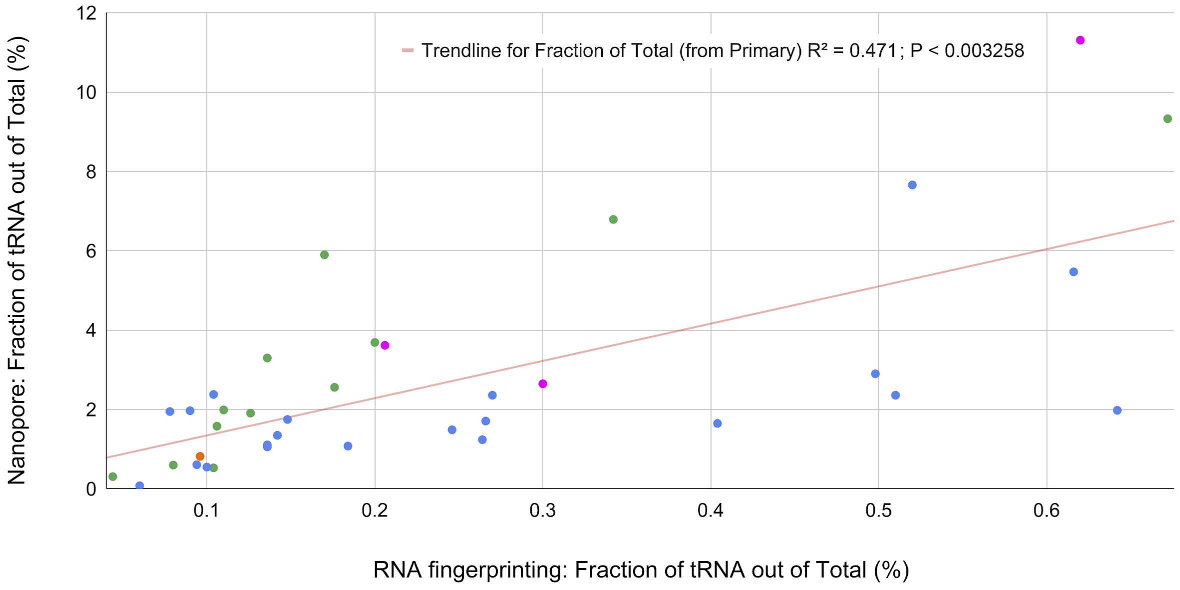


**Supplemental Figure 3. Total tRNA isoacceptor abundance in Nanopore vs RNA fingerprinting.** Each data point represents one tRNA isoacceptor and is colored based on that tRNA's 3' NCCA overhang type: ACCA = blue, CCCA = orange, GCCA = green, and UCCA = pink. The abundance data from RNA fingerprinting[1] has been averaged across five growth phases. There was a moderate positive correlation ($R^2 = 0.471$; $P < 0.0033$).

(1) Dong, H., Nilsson, L. & Kurland, C. G. Co-variation of tRNA abundance and codon usage in Escherichia coli at different growth rates. *J. Mol. Biol.* 260, 649–663 (1996).

A)

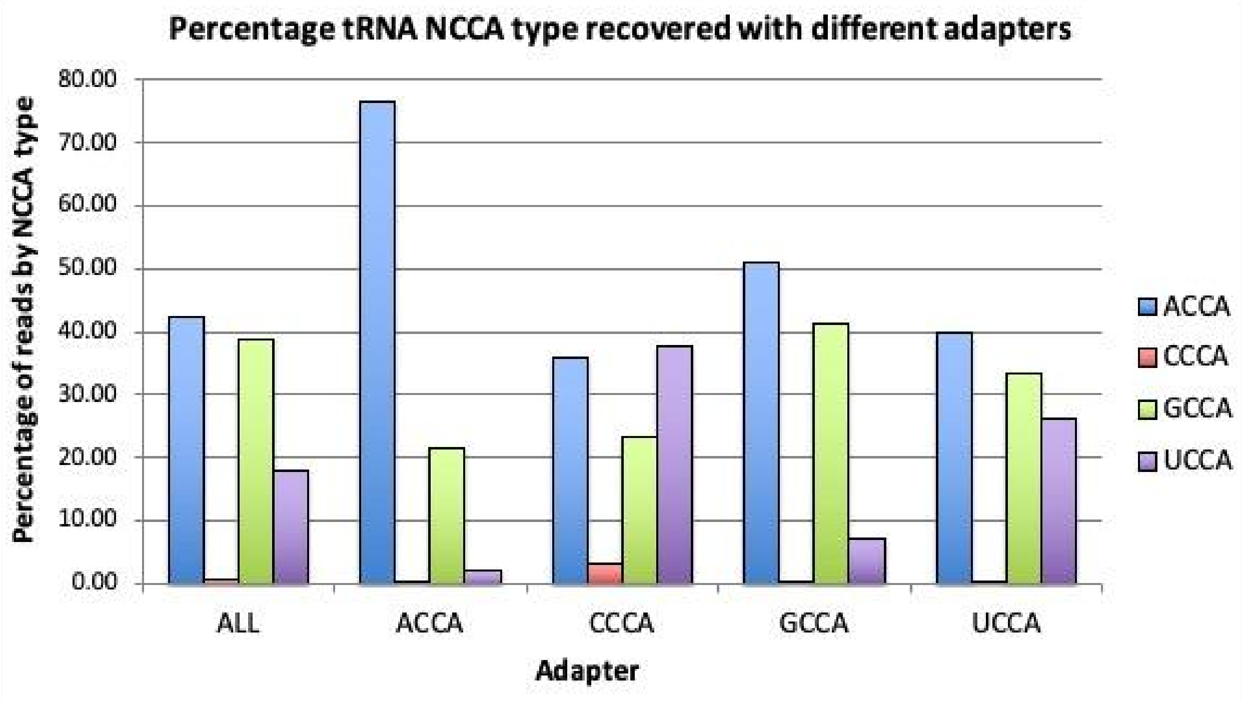


B)

| Percent NCCA type | All Adapters | ACCA Specific Adapter | CCCA Specific Adapter | GCCA Specific Adapter | UCCA Specific Adapter |
|---|---|---|---|---|---|
| ACCA tRNAs | 42.56 | 76.43 | 35.18 | 50.86 | 40.02 |
| CCCA tRNAs | 0.83 | 0.04 | 3.13 | 0.34 | 0.27 |
| GCCA tRNAs | 38.69 | 21.45 | 23.33 | 41.46 | 33.55 |
| UCCA tRNAs | 17.92 | 2.08 | 37.72 | 7.34 | 26.16 |
| Aligned reads MAPQ > 0 | 73,161 | 110,918 | 36,821 | 41,353 | 178,132 |

**Supplemental Figure 4. Percentage of tRNA aligned reads by NCCA termini using all adapters or a single NCCA complementing adapters.** The bar chart shows the percentage of aligned tRNA reads recovered by NCCA type using all adapters or only the adapter specific to ACCA, CCCA, GCCA or UCCA terminating tRNAs (A). In this chart aligned reads for ACCA, CCCA, GCCA and UCCA terminating tRNA are showing in blue, red, green and purple respectively. This data is shown in table form in (B). For each class of tRNA classified by NCCA terminus the percentage of reads is highlighted in the “All adapters” and “Specific Adapter” columns for comparison. In all cases, the percentage of specific NCCA type tRNA recovered increased when the specific adapter was used compared with using all adapters.

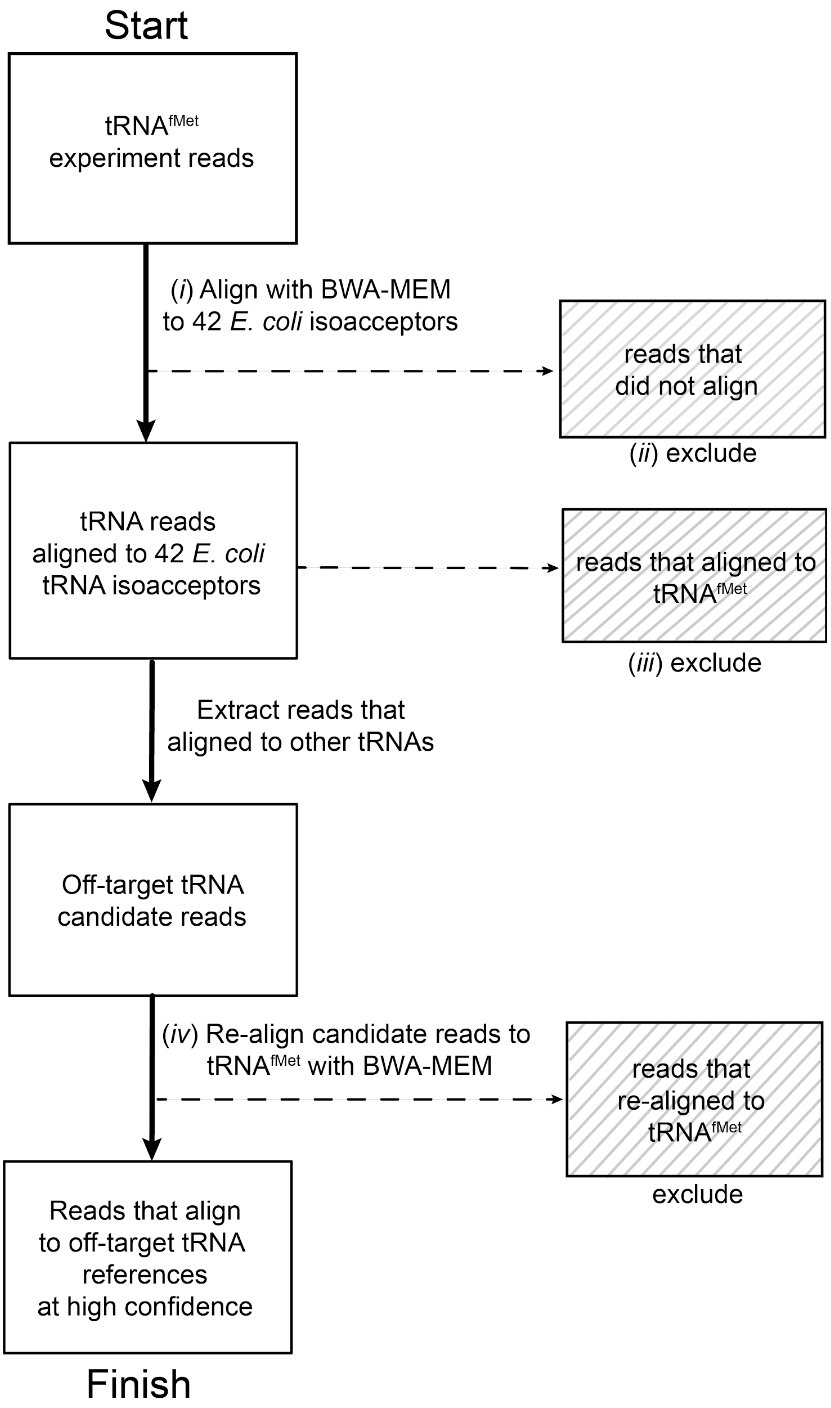


**Supplemental Figure 5. Computational identification of off-target tRNAs in purified samples.** The flowchart below uses a purified $tRNA^{fMet}$ Nanopore experiment as an example. Reads which were systematically excluded are indicated in shaded boxes on the right. *(i)* Nanopore reads for the purified sample were aligned to the curated total tRNA reference. *(ii)* Unaligned reads from *(i)* were excluded. *(iii)* Reads that aligned to $tRNA^{fMet}$ in *(i)* were also excluded. The remaining reads were candidates for off-target tRNAs. *(iv)* These reads were then verified by re-alignment to the $tRNA^{fMet}$ reference. Unaligned reads from *(iv)* were classified as high-confidence off-target tRNAs.

**Supplemental Materials and Methods**

**Generating a canonical tRNA$^{Ala1\ (VGC)}$ control using IVT**

Oligonucleotides for making all canonical tRNA$^{Ala1\ (VGC)}$
Promoter strand (T7 promoter site is underlined)
5′CATCATCATTTAATACGAC3′

Template strand
5′TTTTTTTTTTTTTTTTTACTACCTAAGAGCAAGAAGAAGCCTGGTGGAGCTATGCGGGATCGAACCG
CAGACCTCCTGCGTGCAAAGCAGGCGCTCTCCCAGCTGAGCTATAGCCCCACCAGGCTTCTTCTTGCT
CTTAGGGATGATGATGATGATGATGATGATGATGATGATCTATAGTGAGTCGTATTAAATGATGATG3′

*In vitro* transcription of *E. coli* tRNA$^{Ala1}$
*In vitro* transcription (IVT) was performed using the HiScribe T7 Quick High Yield RNA Synthesis Kit (NEB, E2050). The sequences of the DNA template oligonucleotide and the primer containing the T7 promoter site are shown above. The construct was designed to produce an 180 nt long product. This product has a 61 nt extension 5′ of the tRNA that included the RNA portion of the 5′ splint adapter. This was followed by the tRNA, a 24 nt 3′ extension of including the 3′ splint adapter strand, followed by a 17 nt 3′ polyA tail. The purpose of the 5′ extension was to insure full coverage at the 5′ end of the tRNA. The 3′ polyA tail made the construct compatible with the RTA adapter (ONT) designed for mRNA sequencing(Garalde et al., 2018).

165 pmol of the T7 promoter oligonucleotide (1.65 µl of 100 µM stock) and 33 pmol (0.66 µl of 50 µM stock) of template oligomer were diluted in 10mM Tris-HCl (pH 8.0), 50 mM NaCl and 1 mM EDTA in a total volume of 7 µl. The DNA was hybridized by heating to 75°C for 1 min, then slowly cooling to 23°C. For the IVT reaction, the hybridized oligomers were added to 10 µl NTP mix, 2 µl T7 RNA polymerase and 1 µl (40U/µl) RNAsin Plus (Promega), then incubated for 10 hrs at 37°C. The reaction was DNAse I (RNAse-free)(2,000 units/mL)(NEB) treated for 30 min at 37°C, purified with 1.6X Agencourt RNAClean XP beads (Beckman Coulter), washed with 200 µl 70% ETOH and eluted with 30 µl NF $H_20$. The concentration was determined by nanodrop.

PAGE Gel separation and excision of the IVT product
Six µg of IVT product was diluted to 1X with 2X RNA Loading Solution (NEB). Standard preparation, gel run parameters, staining and excision of full length IVT product (~125nt) were as described for the "PAGE Gel separation and excision of the tRNA/splint ligation product" in the Materials and Methods.

Gel purification by electroelution of the IVT product
The excised IVT product was electroeluted using D-tube dialyzer Midi columns (Novagen) and ethanol precipitated as described in "Gel purification of tRNA/splint ligation product" (Materials and Methods). Following ethanol precipitation, two washes were done by adding 200 µl of freshly made 70% ETOH. The tube(s) were spun for 15 min at 12,000 g in a microfuge and the ethanol from the pellet. After the second wash, the pellets were air dried for 10 min, then resuspended and pooled in a total of 16 µl NF $H_2O$. The concentration was measured with the Qubit HS fluorometer assay or nanodrop.

Library preparation of IVT generated canonical tRNA$^{Ala1}$
350-500 ng of gel purified RNA was used for the library. The IVT generated tRNA was designed to include a 17 nt poly(A) tail, and the standard SQK-RNA002 protocol for direct sequencing of mRNA was followed.

Notes on the sequencing of IVT control

Using this approach, minION sequencing generated 522,342 reads. Alignments of these reads to a reference containing the 42 tRNA isoacceptors gave 452,792 primary alignment to the expected $tRNA^{Ala1}$ (VGC anticodon) and 11,543 reads aligning to the $tRNA^{Ala2}$ (GGC anticodon). The throughput for this IVT-generated tRNA was better than the runs using synthetically generated canonical tRNA that underwent the splint adapter ligation (Supplemental Table 4) and suggests it is a good approach for generating canonical tRNA controls.
